# Transcriptional and functional profiles of muscarinic receptor-expressing neurons in primate lateral prefrontal and anterior cingulate cortices

**DOI:** 10.1101/2025.05.23.655820

**Authors:** Alexandra Tsolias, Chromewell A. Mojica, Raghad Yamani, Sonal D. Khanna, Salam Al Abdullatif, Benjamin J. Snyder, Wayne Chang, Teresa Guillamon-Vivancos, Joseph Goodliffe, Angela L. Capriglione, Isabel Luisa Tan Palanca, Joaquin Martinez, Joshua D. Campbell, Jennifer I. Luebke, Ella Zeldich, Maria Medalla

**Affiliations:** Department of Anatomy & Neurobiology, Boston University Chobanian & Avedisian School of Medicine; Department of Medicine, Boston University Chobanian & Avedisian School of Medicine; Center for Systems Neuroscience, Boston University; Neurophotonics Center, Boston University; Bioinformatics Program, Boston University; Department of Biochemistry, Faculty of Science, King AbdulAziz University, Jeddah. Saudi Arabia

## Abstract

Acetylcholine modulates anterior cingulate (ACC) and lateral prefrontal (LPFC) cortices for cognitive-motivational integration, via specific m1-m4 muscarinic receptors (mAChR) encoded by *CHRM1-4* genes. Single-nucleus RNA sequencing and mRNA-protein histology in macaques revealed *CHRM3* to be the most enriched mAChR gene in neurons, while m1 predominates at the protein level, likely due to nuclear retention of *CHRM3* and cytoplasmic trafficking of *CHRM1. CHRM3* and *CHRM1* showed strong co-expression and functional overlap, and were transcriptomically-distinct from *CHRM2,* which was uniquely enriched in deep layer excitatory and *PVALB*+ inhibitory neurons. Although *CHRM*+ cell distributions were similar between areas, *CHRM1–3*+ excitatory neurons in ACC exhibited upregulation of synaptic plasticity genes relative to LPFC. Functional *in vitro* experiments confirm a more robust cholinergic-mediated decrease in excitatory:inhibitory synaptic ratio in ACC than in LPFC neurons, accompanied by compensatory changes in spine morphology. These findings highlight region-specific acetylcholine signaling essential for flexible processing, learning and memory.

## Introduction

The lateral prefrontal cortex (LPFC) and the anterior cingulate cortex (ACC) are two key regions of the frontal executive control network, essential for higher-order cognitive functions [reviewed in^1, 2^]. Acetylcholine (ACh) is a robust neuromodulator of excitatory and inhibitory circuit dynamics and plasticity within these higher-order areas, mediating learning, memory and flexible behavior^3–7^;reviewed in^8–11^]. The ACC, as part of the limbic system, receives markedly denser cholinergic projections compared to the LPFC^12, 13^. The ACC communicates with limbic centers for arousal and emotions, and sends contextual and motivational information to LPFC to guide behavior^14–24^. In rodent limbic medial temporal lobe structures [reviewed^25^], ACh enables switching across arousal states and oscillatory dynamics to mediate selective gating of inputs for motivational and mnemonic processing^4, 26^. Similar mechanisms may be at play within ACC-LPFC networks that integrate cognitive-motivational information across diverse timescales [^24, 27–32^; reviewed in^11, 33^]. However, effects of cholinergic modulation on distinct cell types in these diverse higher-order frontal areas are largely unknown in primates.

Diverse cortical cholinergic modulation is predominantly mediated by metabotropic G-protein coupled muscarinic receptor (mAChR) m1-m4 subtypes of two broad pharmacological classes, associated with distinct downstream signaling pathways [^34^; reviewed in^9, 10^]. The M1/M3 class includes postsynaptically localized m1 and m3 subtypes coupled to G_q/11_ proteins, which promote activation of protein kinase C (PKC) and downstream neuronal excitation [^34, 35^ reviewed in^10, 36^]. Conversely, the M2/M4 class includes presynaptic m2 and m4 subtypes coupled to G_i/o_ proteins, which inhibit the function of adenylyl cyclase and reduce cAMP^9, 10^. Activation of M2/M4 receptors alters the activity of potassium (K^+^) channels and calcium (Ca^2+^) channels, resulting in the suppression of neurotransmitter release^34, 37, 38^.

The binding of ACh to different mAChRs can result in diverse excitatory or inhibitory effects depending upon the initiation of specific signaling cascades^39^. In previous studies^40, 41^, we found that the laminar distribution and subcellular structural localization of m1 and m2 mAChR proteins are distinct across frontal areas and neuronal subclasses. The current study aims to identify transcriptional regulation associated with mAChR m1-4 subtype gene (*CHRM1-4*) expression in ACC and LPFC excitatory (ExNs) and inhibitory neurons (InNs), using single-nucleus RNA sequencing (snRNAseq), and to assess the functional effects of cholinergic activation on excitatory and inhibitory synapses using the cholinergic agonist, carbachol (CCH), and *in vitro* whole-cell patch-clamp electrophysiological recordings in adult rhesus monkeys. Our findings pave the way to understanding how region-and cell-specific cholinergic modulation may differentially affect distinct cognitive domains, and learning and memory functions, disrupted in cognitive disorders.

## Results

### Cell-specific mAChR subtype gene expression patterns are generalized across ACC and LPFC: predominant enrichment of CHRM3+

For assessments of cell-type specific transcriptional profiles associated with mAChR enrichment, we isolated a total of 9195 nuclei from the ACC area 24, and 8147 nuclei from the LPFC area 46 of adult rhesus monkeys, for snRNA-seq (see Methods, Fig. 1A, Extended Fig. 1A-D). Analyses identified clusters corresponding to 7 major cell types based on expression of canonical cell-specific markers: ExN, InN, microglia, astrocytes (Astr), oligodendrocytes (OL), oligodendrocyte precursor cells (OPCs) and “other” cells (mainly pericytes and endocytes)^42–46^ (Fig. 1B, Extended Figure 1C, E-K, and Table S2).

**Figure 1.**
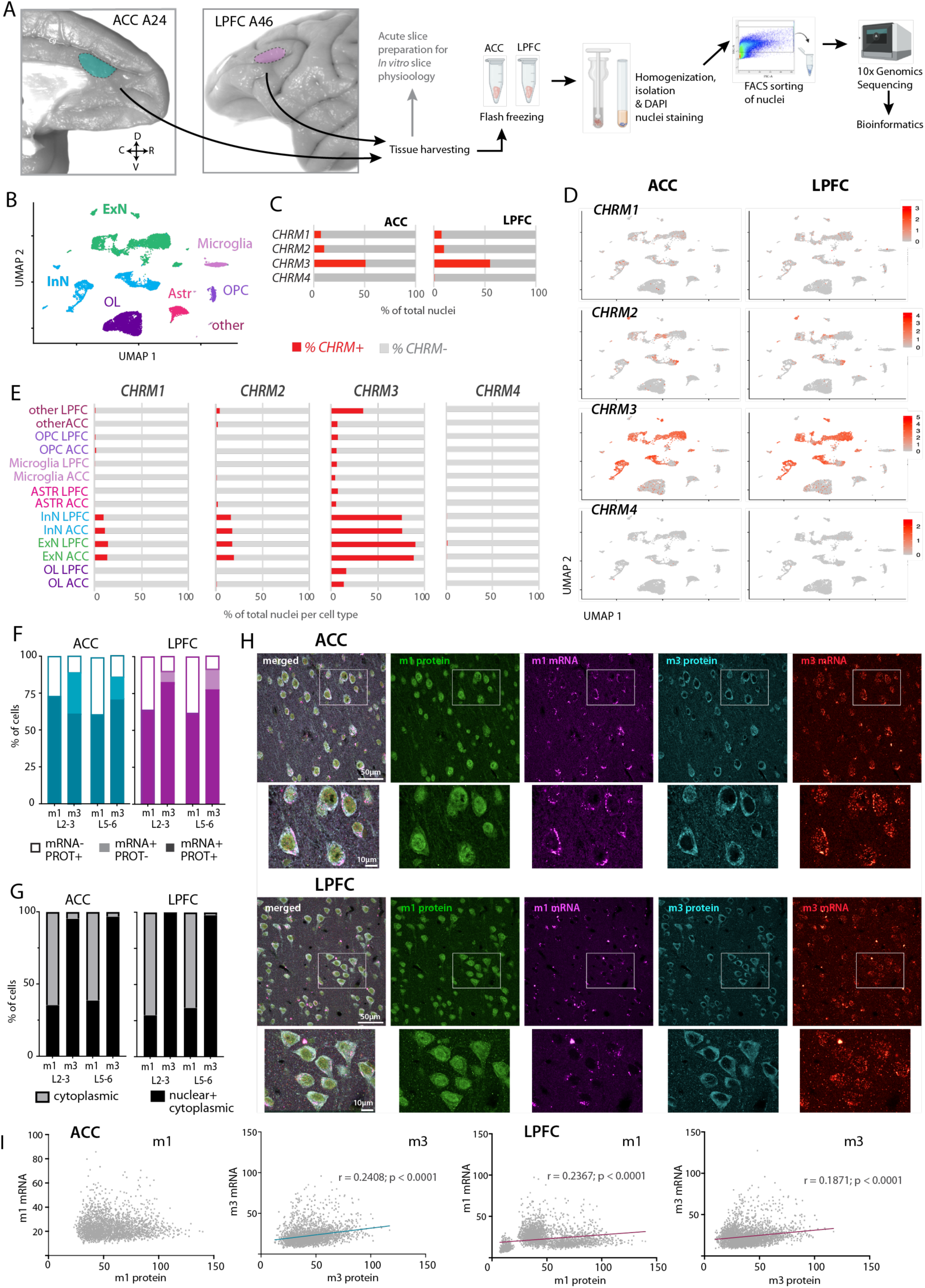
Predominant neuronal expression and nuclear retention of *CHRM3* in ACC and LPFC. (**A**) Photographs of the rhesus monkey brain showing locations of ACC (area 24) and LPFC (area 46) regions of interest and schematic of experimental workflow. (**B**) UMAP plot based on snRNA-seq showing transcriptomically distinct clusters corresponding to major cell types in ACC and LPFC (see Extended Fig. 1). (**C**) Stacked horizontal bar graph showing percentage of total nuclei in ACC and LPFC expressing *CHRM1, CHRM2, CHRM3, CHRM4*; (**D**) UMAP feature plots showing expression pattern of the *CHRM1-4* in ACC and LPFC; (**E**) Proportion of nuclei expressing *CHRM1-4* out of total nuclei by cell type for each area. (**F**) Proportion of total m1+ and total m3+ mRNA and protein expressing cells in ACC and LPFC. Open bars show the proportion of cells that are mRNA negative but protein positive (mRNA-/PROT+), and filled bars show the proportion positive for mRNA with (mRNA+/PROT+) or without (mRNA+/PROT-) protein. Note the presence of m3+ cells expressing mRNA but no protein (mRNA+/PROT-). (**G**) Proportion of cells showing mRNA label localized only in the cytoplasm, or within nuclei and cytoplasm. Nuclear vs cytoplastic localization was based on the central versus peripheral localization of label and its overlap with MAP2 cytoplasmic labeling. (**H**) Representative z-maximum projection confocal images (5 optical stacks) from ACC and LPFC L2-3 showing the cellular/ subcellular distribution of m1 protein (green) and mRNA (magenta) and m3 protein (cyan) and mRNA (red). Note the predominantly cytoplasmic localization of *CHRM1*, versus the diffused distribution of *CHRM3* within nuclear and cytoplasmic compartments. (**I**) Scatter plot of mRNA/protein density within individual cells and linear regression analyses in ACC and LPFC, showing correlations of m1 and m3 mRNA vs protein.

Distinct neuronal and glial cell clusters had detectable expression of the *CHRM1-4* genes for m1-m4 mAChR subtypes, in patterns that are remarkably similar in ACC and LPFC (Figure 1C-E). Unexpectedly, while m1 and m2 mAChRs predominate at the protein level in the cortex^47^, our findings revealed *CHRM3* (m3) to be the most widely expressed mAChR mRNA in both ACC and LPFC, which was expressed on ∼51-55% of all cells (Fig. 1C-E). Only about ∼7% and ∼10% of all cells expressed *CHRM1* (m1) and *CHRM2* (m2), and a very small proportion of ∼0.5-0.7% (< 100 nuclei) of all cell types expressed *CHRM4* (m4).

Examination of mAChR gene enrichment within each major cell type revealed that *CHRM3*+ neurons comprise 89-91% of total ExN and ∼76% of total InN in both areas (Fig. 1E). In contrast, only 10-14% and 16-20% of all neurons expressed *CHRM1* and *CHRM2*, respectively (Fig. 1E). *CHRM1* and *CHRM2* were barely detectable in a small proportion of non-neuronal cells (0–2% for *CHRM1* and 0-4% for *CHRM2*). *CHRM4* was exclusively expressed in neurons. In contrast, *CHRM3* was expressed in 14-17% of total oligodendrocytes, 6-8% of total OPCs, 6-8% of total astrocytes and 5-7% of total microglia (Fig. 1E).

Within each *CHRM*-expressing subpopulation, > 90% of the *CHRM*+ cells were neurons, with ∼*61-82%* ExNs and 18-33% InNs (Extended Fig. 1K). *CHRM3* had the highest proportion expressed in the non-neuronal cells, specifically in oligodendrocytes (7.5%) and astrocytes (1.05%) (vs *CHRM1/2* cells consisting of ∼1% oligodendrocytes and 0.3-0.4% astrocytes; and no expression within *CHRM4*; Extended Fig. 1K). Thus, neuronal versus non-neuronal expression of *CHRM* was dependent on the specific *CHRM* gene subtype.

### Predominant nuclear mRNA localization and mRNA:protein correlation in m3+ expressing neurons

The widespread expression of *CHRM3+* compared to other mAChR subtype genes did not align with previous studies at the protein level that showed that m1 is the most predominantly expressed mAChR subtype in the cortex^47^. Thus, histological validation of m1 and m3 mRNA-protein expression was employed using combined IHC and fluorescence in situ hybridization (FISH, see Methods and Supplementary Table S3; Fig. 1F-H). Among all m1 protein+ expressing neurons, ∼61-73% in ACC and 62-64% in LPFC also expressed *CHRM1+* mRNA. Conversely, 27-39% in ACC and 36-38% in LPFC of these m1+ neurons expressed m1 protein but not *CHRM1* (Fig. 1F, H). Almost all m3+ protein expressing neurons (∼ 88-95% in ACC and 95-96% in LPFC) also expressed *CHRM3*. In contrast to m1+ neurons, we found a proportion of neurons with detectable *CHRM3* that were negative for m3 protein (ACC: L2-3 = 29%, L5-6 = 15%; LPFC: L2-3 = 17%, L5-6 = 19%; Fig. 1F, H).

In addition to differences in protein and mRNA co-expression, our analyses revealed a marked difference in intracellular localization of *CHRM1* vs. *CHRM3*. Most *CHRM1* was localized to the cytoplasm, with weak or no labeling in the nucleus (% cytoplasmic, ACC: 61-64%; LPFC: 66-71% of m1mRNA+ cells; Fig. 1G, H). In contrast, *CHRM3* was detected in both the nucleus and cytoplasm of all cells (% nuclear + cytoplasmic, ACC: 95-97%; LPFC: 98-100% of m3mRNA+ cells). These data suggest differences in the mechanisms regulating mRNA export and translation between the subtypes of muscarinic receptors.

We therefore assessed the correlation between protein and mRNA levels in m1 vs m3 expressing neurons (Fig. 1I). For m3+ cells in both ACC and LPFC, we found a significant positive correlation between mRNA and protein particle density (ACC: Pearson’s r = 0.2408; LPFC: r = 0.1871; p < 0.0001). For m1+ cells, mRNA and protein expression was only correlated in LPFC (r = 0.23567; p < 0.0001) but not in ACC (p = 0.3792). These relationships further support the region-specific differential mRNA and protein dynamics in m1 vs m3 neurons, with mRNA and protein being more strongly correlated in m3 compared to m1 expressing neurons.

### CHRM1-4 mRNA was widely co-expressed in patterns that differed across layer-specific ExN subpopulations

We assessed the relative distribution of ExNs expressing/co-expressing *CHRM1-4* and found a similar pattern between the ACC and LPFC. Within both areas, the majority (∼60%) of ExNs expressed *CHRM3* only (Fig. 2A, dark blue). The second largest population, consisting of 27-28% of all ExN, were those co-expressing two or more mAChRs (Fig. 2A, dark grey). The rest of the ExN population was comprised of ∼4% *CHRM2+* only, and 5-7% with no mAChR gene expression (Fig. 2A, orange and light grey, respectively). We found no ExNs that exclusively expressed *CHRM4*, as it was always co-expressed with another mAChR gene.

**Figure 2.**
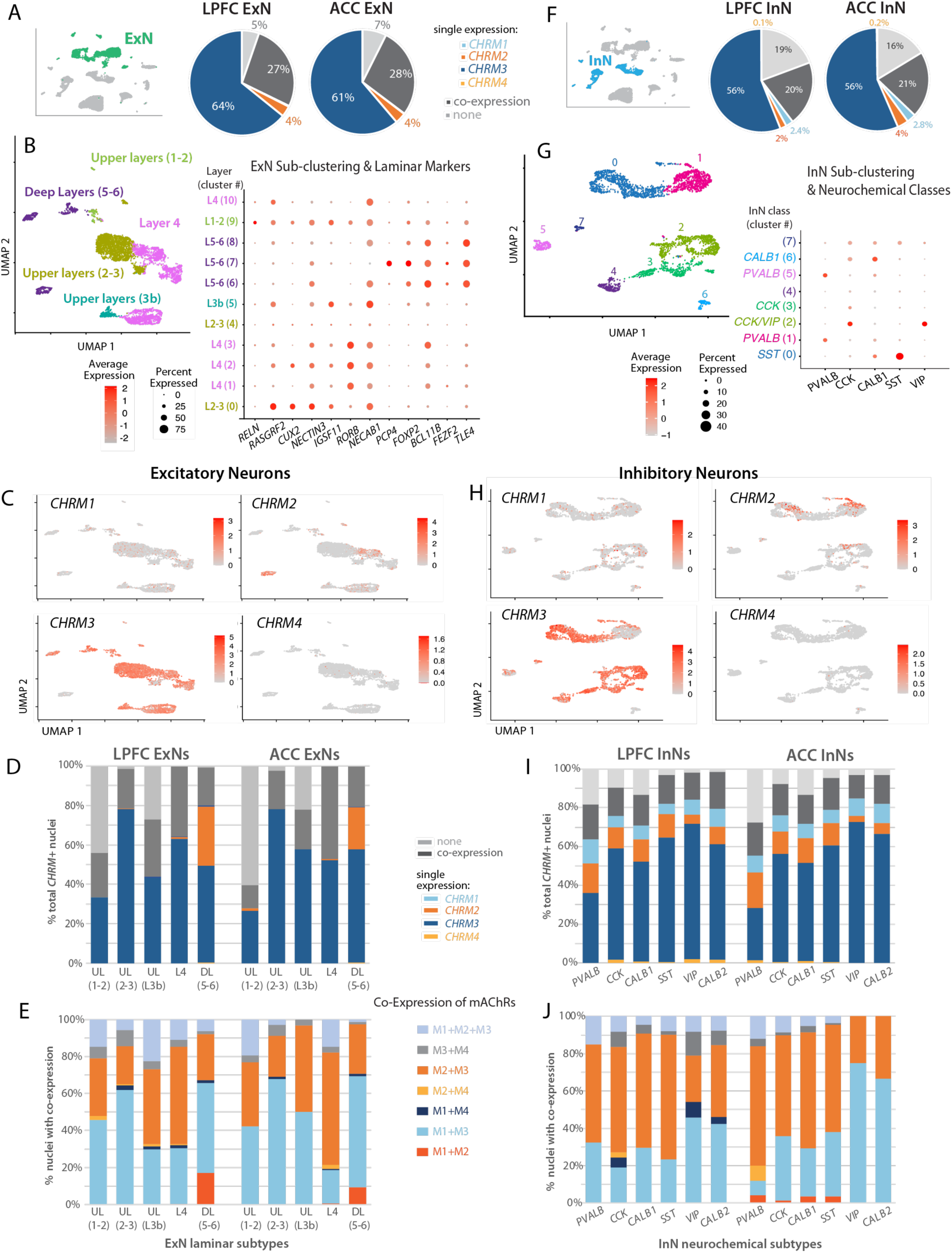
*CHRM1-3* show distinct distribution across layer-specific ExNs and neurochemical InN subclasses in ACC and LPFC. (**A)** Pie chart showing the overall proportion of ExN expressing/co-expressing specific *CHRMs* in ACC and LPFC. Inset shows UMAP highlighting the population of ExNs (green) together with other cell types. (**B**) UMAP plot (left) after re-clustering of ExNs annotated based on layer-specific subpopulations identified via expression of layer-specific genes in each cluster shown in dot plot (right; see Extended Fig. 2 and Table S2). (**C**) Feature plots showing the expression pattern of *CHRM1-4* following the re-clustering of ExNs. (**D**) Within each ExN laminar subpopulation: proportion of cells with single, coexpression (>1) or no expression of *CHRM1-4* are presented as a stacked bar plot, UL=upper layers, DL=deep layers. (**E**) Within ExNs with coexpression, proportion of cells with specific combinations of *CHRM1-4* coexpression are presented as a stacked bar plot. (**F**) Pie chart showing the overall proportion of InN expressing/co-expressing *CHRMs* in ACC and LPFC. Inset shows UMAP highlighting the population of InNs (blue) together with other cell types. (**G**) UMAP plot (left) after re-clustering of InNs showing 7 distinct clusters/subpopulations annotated based on expression of neurochemical markers. Dot plot shows expression pattern of a subset of markers for distinct neurochemical subclasses of InNs based on the literature (see Extended Fig. 2). (**H**) Feature plots showing the expression pattern of *CHRM1-4* following the reclustering of InNs. (**I**) Within each InN subclass, proportion of cells with single, coexpression (>1) or no expression of mAChR genes are presented as a stacked bar plot. (**J**) Within InNs with coexpression, proportion of cells with specific combinations of *CHRM1-4* coexpression coexpression.

More granular transcriptomic analysis of ExN subpopulations via re-clustering of cell types annotated as ExN using Algorithm and resolution, revealed differences in *CHRM1-4* expression/co-expression patterns associated with expression of layer-specific genes. The re-clustering analysis yielded 11 clusters of ExNs (Fig. 2A, B, Extended Fig. 2A-E) distributed and annotated based on distinct expression of layer-specific genes^48–51^ (Fig. 2B, C, Supplementary Table S2). For the largest subpopulation of upper layer (UL) 2-3 ExNs, ∼80% of neurons expressed only *CHRM3*, and ∼20% co-expressed two or more mAChR*s* (Fig. 2D, dark blue). Layer 4 ExN cluster expressed either only *CHRM3* or co-expressed mAChRs, with a more equal distribution (60-40% ratio). Similarly, 45-60% of UL3b ExNs expressed only *CHRM3* and the rest either co-expressed >2 mAChR*s (20-30%)* or did not express any mAChR (20-25% of ExN). While the deep layer (DL) L5-6 ExN cluster also included ∼50-60% ExNs expressing only *CHRM3,* this subcluster was uniquely enriched with ExNs expressing only *CHRM2* (20-30% of DL5-6 ExN, sub-cluster 7; Fig. 2D, orange). In contrast to other clusters, the UL1-2 ExN cluster had a relative enrichment of cells (∼40-60%) that did not express any mAChR (of total UL1-2 ExNs).

Further analyses of the specific combinations of *CHRM1-4* co-expression revealed layer-specific ExN distribution patterns among subpopulations (Fig. 2E). The majority of ExNs in UL1-2, UL2-3, L4 and DL5-6 clusters in both areas, co-expressed *CHRM1*+/*CHRM*3+ (Fig. 2E, light blue). In contrast, ExN in UL3b and L4 clusters with mAChR co-expression, were predominantly *CHRM2+/CHRM*3+ (Fig. 2E, orange). Interestingly, DL5-6 ExNs in both areas had a relatively higher proportion of cells co-expressing *CHRM1+*/*CHRM*2+ as compared to other clusters (2-3% in DL5-6 vs < 0.5%; Fig. 2E, dark orange). Within all ExN subpopulations, < 5% co-expressed *CHRM4* with *CHRM1-3*. These results show distinct patterns of mAChRs mRNA enrichment across layer-specific ExN subtypes.

### Patterns of mAChR subtype mRNA expression/co-expression distinguished two broad groups of InN subpopulations

Similar to ExN, most (∼56%) of the InN in ACC and LPFC expressed only *CHRM3* (Fig. 2F, dark blue), while the rest (∼19-21%) co-expressed two or more mAChRs (Fig. 2F, dark grey) or had no mAChR expression (Fig. 2F, light grey). A very small proportion of InN were either only *CHRM2+* (2% in LPFC vs 4% of ACC) or *CHRM*1+ (2.4% in LPFC vs 2.8% of ACC; Fig. 2F). Interestingly, in contrast to ExNs, we found < 0.5% of InNs exclusively expressing *CHRM4*. Analyses of InN subpopulations via re-clustering revealed 8 subclusters of InN that were distributed and annotated based on the expression of neurochemically distinct InN genes ^52–56^ (Fig. 2G, H; Extended Fig. 2E-H). Specifically, parvalbumin positive (*PVALB+*), somatostatin (*SST+*), vasoactive intestinal peptide (*VIP+*) neurons formed distinct clusters (Fig. 2G). A subset of *SST*+ neurons (cluster 7) co-expressed calbindin (C*ALB1*), *VIP,* and cholecystokinin genes (*CCK*; Fig. 2G).

Our data revealed that the patterns of mAChR expression/co-expression differentiated neurochemically-distinct InNs cell types. Specifically, for all neurochemical subclusters, except *PVALB+,* > 50% of InN consisted of only *CHRM3+*. *PVALB+* InN cluster had the lowest proportion of only *CHRM3+* (30-35% of *PVALB+* InN; Fig. 2I, dark blue) and the highest expression of only *CHRM2+* (15%, Fig. 2I, orange). Further, our dataset revealed two distinct mAChR co-expression patterns, distinguishing two major groups of InNs (Fig. 2J). The first group consisted of *PVALB+, CCK+, CALB1+,* and *SST+* InNs, which mainly co-expressed *CHRM2+/CHRM3+* (54-64% in ACC; 53-67% in LPFC; Fig. 2J, orange). In contrast, the second group, consisted of *VIP+* and *CALB2+* InNs, predominantly co-expressing *CHRM1+/CHRM3+*, and these InNs were more enriched in ACC than in LPFC (67% in ACC vs 42% in LPFC of *VIP+*; and 75% in ACC vs 45% in LPFC of *CALB2+* InNs; Fig. 2J, light blue).

### Differentially expressed genes (DEGs) across CHRM1-3 enriched neurons show functional overlap of m1 and m3, and their distinction from m2

To assess whether mAChR gene expression was correlated with distinct transcriptomic profiles related to receptor-specific downstream effectors, we identified DEGs cross *CHRM1-3* enriched ExN and InN subpopulations within each region. For this purpose, we performed pseudobulk analysis of the total ExN and total InN cell types and identified DEGs for the following comparisons: *CHRM1+* vs. *CHRM2+; CHRM3+* vs. *CHRM2+,* and *CHRM1+* vs. *CHRM3+ (*Fig. 3A; Extended Fig. 3A, B). Overall, the data point to the similar transcriptomic profiles of *CHRM1+* and *CHRM3+* neurons, which were both distinct from *CHRM2+* neurons. *CHRM1+ vs*.

**Figure 3.**
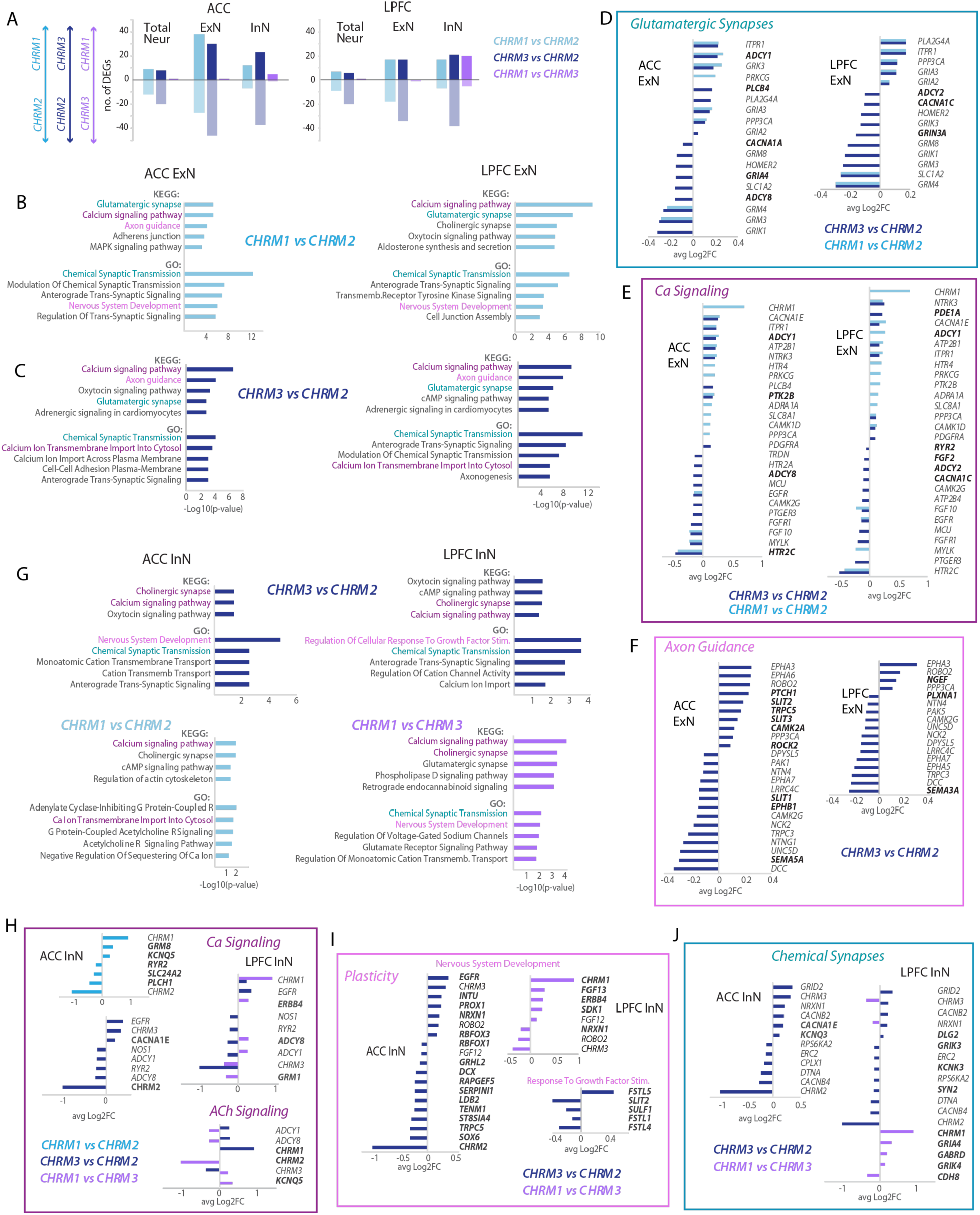
*CHRM* enrichment define cell-specific transcriptomic signatures aligning with mAChR functional classes. (**A**) Bar graph of the number of DEGs in total, ExN or InN enriched in *CHRM1* relative to *CHRM2* (light blue); enriched in *CHRM3* relative to *CHRM2* (dark blue); and enriched in *CHRM1* relative to *CHRM3* (purple). (**B**) Top significantly enriched KEGG pathway (top bars) and GO biological processes (bottom bars) terms generated based on DEGs between *CHRM1+* relative to *CHRM2+* ExNs (light blue) and (**C**) *CHRM3+* relative to *CHRM2+* ExNs (dark blue) The data was generated using Enrich and significance enrichment was defined as p-value<0.05, Benjamini. (**D-F**) Fold-change of top *CHRM1* vs. *CHRM2* and *CHRM3* vs. *CHRM2* DEGs within ACC and LPFC ExNs, related to: (**D**) KEGG term “Glutamatergic Synapse” and GO “Chemical Synaptic Transmission GO: (teal in panels B and C). (**E**) KEGG term “Calcium Signaling”, and GO term “Calcium transmembrane import into the cyosol” GO: (purple in panels B and C) (**F**) KEGG term “Axon Guidance”, and GO term “Nervous System Development”; (**G**) KEGG pathway and GO terms from *DEGs* between *CHRM3+* relative to *CHRM2+* InNs (dark blue), *CHRM1+* relative to *CHRM2+* InNs (light blue), and *CHRM1*+ relative to *CHRM3+* InNs (purple). (**H-J**) Fold-change of top *CHRM1* vs. *CHRM2, CHRM3* vs. *CHRM2,* and *CHRM1* vs. *CHRM3* DEGs within ACC and LPFC InNs, related to: (**H**) KEGG/GO term “Ca Signaling” and “Ca Ion Transport”, and KEGG term “Cholinergic Synapse”; (**I**) Plasticity related terms – GO Nervous system development GO and Regulation of Growth Factor Stimulation GO; (**J**) GO term “Chemical Synaptic Transmission GO. See Supplementary Tables S4 and S5 for a full list of enriched terms and DEGs, and Extended Fig. 3 for volcano plots.

*CHRM2+,* and *CHRM3+ vs. CHRM2+* comparisons resulted in a high number of DEGs genes in both ACC and LPFC ExNs and InNs (Fig. 3A). In contrast, *CHRM1+* vs. *CHRM3+* yielded almost no DEGs in ExNs (1 DEG for each area), a very a low number of DEGs in InNs, (ACC 6, LPFC 25; Fig. 3A, Supplementary Table S5). Interestingly, the degree of similarity between *CHRM1+* and *CHRM3+* seem to be more pronounced for ExNs compared to InNs. For InNs, *CHRM3+* vs. *CHRM2+* resulted in a higher number of DEGs as compared to *CHRM1+* vs*. CHRM2+* for both areas (ACC 60 vs. 19; LPFC 59 vs. 24 DEGs, respectively; Fig. 3A), suggesting receptor-specific differences between *CHRM1+* and *CHRM3+*. Further, the ExN populations revealed region-specific distinctions in the number of DEGs. In the ACC ExNs, *CHRM1+* vs*. CHRM2+*, and *CHRM3+* vs*. CHRM2+* yielded a comparable number of DEGs (65 vs. 76 DEGs, respectively Fig. 2A). However, in LPFC ExNs, the number of DEGs was slightly higher for *CHRM3+* vs*. CHRM2+* compared to *CHRM1+* vs*. CHRM2+* (51 vs. 35 DEGs, respectively).

To evaluate how distinct transcriptomic profiles of mAChR expressing cells relate to function, we utilized functional annotation analyses using KEGG pathway^57, 58^, and gene ontology (GO) analysis^59, 60^ in Enrichr^61–63^. For ExNs, Enrichr analysis of DEGs from *CHRM1+* vs*. CHRM2+* and *CHRM3+* vs*. CHRM2+* comparisons yielded similar significantly enriched (p < 0.05) KEGG pathway and GO terms for biological processes in both areas (Fig. 3B-F, Supplementary Tables S5 and S6). The top enriched terms had overlapping gene sets related to four major functions: synaptic transmission (e.g. Glutamatergic Synapses, Chemical Synaptic Transmission, Synapse Organization; Fig. 3B-D); calcium signaling (Ca Signaling, Ca Ion Transport, Fig. 3B,C,E), axon guidance/plasticity, and development (Axonogenesis, Neuron Projection Guidance, Cell adhesion, MAPK2 and RAP1 pathways that regulate cell polarity and growth, Fig. 3B,C,F)^64, 65^. Significantly enriched terms were also related to cAMP/cGMP, G-protein coupled signaling (Fig. 3B-F), consistent with the opposite physiologic effects of the m1/m3 vs m2 receptors on second messenger cascades^66–68^.

These enriched functional terms common to ACC and LPFC were derived from overlapping *CHRM1+/3+* vs*. CHRM2+* ExN DEGs, which include genes for glutamatergic receptors, Calcium and K+ channels, and related to second messenger cascades (Fig. 3D, E). The top *CHRM3+* genes (relative to *CHRM2+)* in ExNs include *GRIA2 and GRIA3 (*ionotropic glutamate receptor genes), *CACNA1E (*R-type Ca Channel) *ADCY1* adenyl cyclase 1, while some of the top *CHRM2*+enriched genes include *HTR2C* (serotonergic receptor), GIRK1 (inwardly-rectifying K+ channel), and *GRM3* and *GRM4* (metabotropic glutamate receptors). In both areas, *CHRM2-* enrichment in ExNs was associated with positive regulators of synapse maturation and stability genes *(UNC5D, EPHA5/7*^69^*, SEMA3A* ^70^*, DCC* ^71^*, TRPC3* ^72^).

In addition to gene sets that were consistent between areas, there was a small subset of *CHRM3+* vs. *CHRM2+* DEGs encoding for adenyl cyclases, glutamatergic receptors and calcium channels, which were unique within ACC and LPFC ExNs (Fig. 3D, E). For instance, two distinct differentially expressed adenyl cyclases were associated with each region: In ACC, *ADCY1* the subtype responsive to Ca Calmodulin^73^, was enriched in *CHRM3+* ExNs, while in LPFC *ADCY2* was enriched in *CHRM2+* ExNs (Fig. 3E). For Ca channels, ACC *CACNA1A*, which encodes for a P/Q type Ca channel concentrated in presynaptic terminals, and LPFC *CACNA1C*, which encodes for an L type calcium channel in somatodendritic domains, were both enriched in *CHRM2+* ExNs (Fig. 3D, E). In addition, *CHRM3+* vs*. CHRM2+* DEGs related to axon guidance, adhesion and plasticity, showed a relatively high proportion in ACC compared to LPFC (Fig. 3F). Specifically, ACC ExNs had a higher representation of *CHRM3+* vs*. CHRM2+* DEGs linked to the ROBO pathways, mediating axon guidance and adhesion, as compared to LPFC (Fig. 3F, *ROBO1* and *SLIT1/2/3*, *SEMA5A*)^74–76^.

Similar to ExNs, functional annotation analyses of DEGs associated with *CHRM1-3* expression in InNs within each area yielded significantly enriched terms related to Ca signaling, chemical synaptic transmission and plasticity (Fig. 3G-J). While the number of DEGs across *CHRM1-3* expressing InNs was smaller in comparison to ExNs, the fold change for each gene was greater, and almost always included the defining *CHRM* in each term. Similar to ExNs, the number of *CHRM3+* vs*. CHRM2+* InN DEGs related to plasticity was greater in ACC than in LPFC, which included genes associated with presynaptic specialization assembly (*ROBO2, TNEM1*)^74–77^ and RNA dynamics important for synapse development (*RBFOX1/3, SOX6*)^78, 79^.

The analyses showed unique DEGs yielded by *CHRM1+* vs*. CHRM2*+ and *CHRM3+* vs*. CHRM2*+ comparisons, especially in LPFC InNs. For instance, the term ‘chemical synaptic transmission’ included DEGs related to GABAergic and Glutamatergic receptors in LPFC *CHRM1+* vs*. CHRM3+* InNs, but related to Ca Channels, K channels and synapse structure in LPFC *CHRM3+* vs*. CHRM2+* InNs (Fig. 3J, Supplementary Table S4, S5). These data further support the transcriptomic heterogeneity of *CHRM1+* vs*. CHRM3*+ InNs, driven primarily by neurotransmitter specificity, that was not found for ExNs.

Taken together these data indicate that distinct *CHRM* enrichment within both ACC and LPFC correlates with transcriptomic signatures linked to key signaling cascades, in patterns consistent with the functional overlap between m1 and m3 mAChRs that are distinct from m2. Further, the degree of this m1 and m3 functional overlap is more pronounced in ExNs than in InNs.

### Greater number of DEGs between ACC and LPFC for CHRM1-3 enriched ExN than for InNs

Within each *CHRM*+ neuron subpopulation, between-region differential gene expression analyses revealed that ExNs exhibited greater regional differences compared to InNs. We identified higher numbers of between-region DEGs for *CHRM1+*, *CHRM2+*, and *CHRM*3+ ExNs (140, 93, and 204 DEGs, respectively) compared to *CHRM2*+, and *CHRM3+* InNs (no DEGs for *CHRM1+*, 8 for *CHRM*2+, and 37 for *CHRM3+*; Fig. 4A-C, see Supplementary Table S6). Enrichr analysis based on between-region DEGs revealed significant enrichment in KEGG pathways and GO terms that were markedly similar between *CHRM1+* and *CHRM3+* ExNs. The top common terms enriched in ACC were related to synaptic plasticity (axon guidance, cell-to-cell adhesion and chemical synaptic transmission; Fig 4D-F). ACC-upregulated DEGs associated with axon guidance and cell-to-cell adhesion, included genes coding for the ROBO pathway members (*ROBO1/2, SLIT3, SEMA3A*), Cadherins (*CDH6/4/9/10/13*), Teneurins (*TNEM1/2/3*), Ephrin signaling components (*EPHA6/7/3,*) and some other scaffolding and adhesion proteins (*NLGN1, DCC, UNC5D)*; Fig. 4G, H) for synapse formation^69, 70, 74–77, 80^. Related between-area DEGs were associated with chemical synaptic transmission (Fig. 4D, d inset, 4F, f inset, and 4I), which included glutamate receptors (*GRIA1/4, GRIN2B/3A, GRID2, GLRA2*), GABA receptors (*GABRA3/5/2*), synaptic structural and adhesion proteins (*NLGN1, ERC1*) and Ca channels (*CACNA1E/B2*). In addition*, CHRM1+* ExN between-region DEGs showed the unique enrichment for RAP1 signaling and Long-Term Potentiation (LTP) terms that are important for synapse plasticity^81–84^ (Fig. 4D). Between-region DEGs in *CHRM1+/3+* ExNs which were associated with synaptic function and plasticity were primarily upregulated in ACC compared to LPFC (Fig. 4D, d inset, 4F, F inset, and 4I). Interestingly, we found that most of the ionotropic GLURs genes were ACC-enriched (relative to LPFC), while GABAergic receptor genes had mixed trends. For *CHRM3+* ExNs, *GABRA3/5/2* and *GABBR1* genes were ACC-enriched, but *GABRB2, GABBR2 and GABRAG3* were enriched in LPFC (Fig. 4F, f inset).

**Figure 4.**
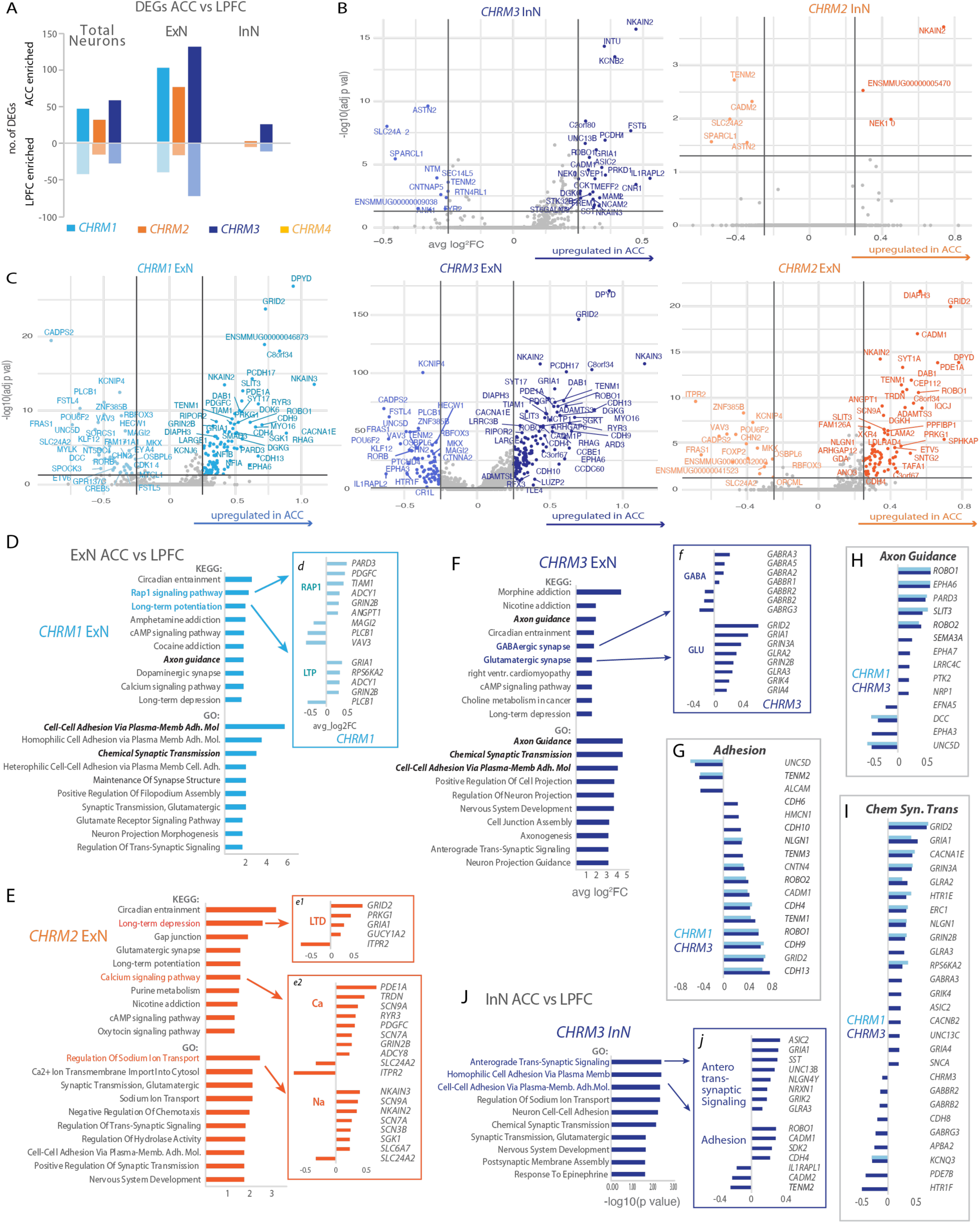
Regional differences in plasticity-related transcriptomic signatures within *CHRM1-3* enriched neurons. (**A**) Bar graph of the number of DEGs enriched in ACC relative to LPFC, within *CHRM1-4+* expressing neurons (total, ExNs, and InNs). (**B, C**) Volcano plots of the DEGs between ACC and LPFC, within *CHRM1-3+* InNs (**B**) and ExNs (**C**). (**D-F**) Top significantly enriched KEGG pathway (top bars) and GO biological processes (bottom bars) terms generated based on DEGs between ACC relative to LPFC for: (**D**) *CHRM1+ ExNs* (light blue). Inset (d) shows fold-change of top ACC vs. LPFC DEGs related to: “Rap1 Signaling” and “Long Term Potentiation”; (**E**) *CHRM2+ ExNs* (orange). Insets (e1, e2) show fold-change of top ACC vs. LPFC DEGs related to “Long Term Depression” (LTD), “Calcium Signaling” (Ca), and “Regulation of Sodium Ion Transport” (Na); (**F**) *CHRM3+* ExNs (dark blue). Inset (f) shows fold-change of top ACC vs. LPFC DEGs related to “GABAergic Synapse” (GABA) and “Glutamatergic Synapse” (GLU). (**G-I**) Fold-change of top ACC vs. LPFC DEGs within *CHRM1* and *CHRM3* ExNs, related to the following common terms (in bold black letters in D, F): (**G**) “Cell-Cell Adhesion via Plasma Membrane Adhesion Molecules” (Adhesion); (**H**) “Axon Guidance”; and (**I**) “Chemical Synaptic Transmission”. (**J**) Top significantly enriched GO biological processes terms generated based on DEGs between ACC relative to LPFC in *CHRM3+* InNs (dark blue). Inset (j) shows fold-change of top ACC vs. LPFC DEGs related to “Anterograde Trans-Synaptic Signaling” and “Homophilic Cell Adhesion Via Plasma Membrane” (Adhesion). The data was generated using Enrichr and significance enrichment was defined as p-value<0.05, Benjamini. See Supplementary Table S6 for a full list of enriched terms and DEGs.

Similar to *CHRM*3+/*CHRM1+* ExNs, a subset of significantly enriched KEGG pathway and GO terms found for *CHRM2+* ExN ACC vs LPFC DEGs were related to Ca signaling, glutamatergic signaling, oxytocin signaling, nicotine, cAMP, adhesion, and development (Fig. 4E). However, we found terms unique to *CHRM2+* ExNs related to Sodium Ion transport and Long-Term Depression (LTD) (Fig. 4E). These DEGs were mostly ACC-enriched relative to LPFC, which include *PDEA1* gene for Phosphodiesterase 1 modulator of cAMP/cGMP activity, the Na channel gene *SCN9A* and cGMP-dependent protein kinase gene, *PRKG1*, which are implicated in nociception^85, 86^ (Fig. 4E, e1, e2, insets). Importantly, *GRID2* and *GRIA1* genes for ionotropic GLUR delta type subunit 2 and GLUR AMPA type subunit 2, respectively, were consistently enriched in ACC relative to LPFC across *CHRM+* expressing ExNs (Fig. 4D-I).

In contrast to ExNs, the comparison between ACC vs. LPFC among the *CHRM+* InN population yielded a markedly lower number of DEGs, with significantly enriched GO terms only for *CHRM*3+. Similarly to ExNs, we found a significant enrichment in the processes related to cell-cell adhesion and trans-synaptic signaling for between-region DEGs in *CHRM*3+ InNs. (Fig. 4J). The genes upregulated in ACC (relative to LPFC) included Cadherins (*CDH4*), genes coding for synaptic adhesion (*NLGN4Y, NRXN1, CADM1*), and presynaptic (*UNC13B*) and postsynaptic (*GRIA1, GIRK2, GLRA3*) specializations (Fig. 4J, j inset; Supplementary Table S6). These findings reveal region and cell-type specific transcriptomic changes as they pertain to *CHRM1-3* enrichment.

### Functional experiments demonstrated differential effects of CCH on synaptic properties in ACC *and LPFC*

Our transcriptomic profiling revealed changes in genes related to synapse stability that are upregulated in ACC relative to LPFC. This led us to investigate differences in synaptic properties between the two areas. Therefore, we assessed the effects the cholinergic agonist, carbachol (CCH), on synaptic currents and structural measures of synaptic plasticity in layer 3 (L3) pyramidal neurons in ACC and LPFC using whole-cell patch clamp recording and intracellular filling. Recordings of spontaneous excitatory and inhibitory post-synaptic currents (EPSC and IPSC) were obtained from L3 pyramidal neurons of ACC (n=12 EPSC, n=11 IPSC) and LPFC (n=11 EPSC, n=9 IPSC; Fig. 5A-D), following 6-8min bath exposure to CCH (10µM, CCH; Fig. 5D and Extended Fig. 5A, B). Most cells in both frontal areas exhibited a decrease in the frequency of EPSCs (77% decrease in ACC, 73% decrease in LPFC) while a smaller proportion exhibited an increase (15% ACC, 27% LPFC) or no change in frequency (8% ACC) (Fig. 5F). A comparison of the sub-population of neurons that exhibited decreased EPSC frequency found that CCH significantly decreased the mean EPSC frequency only in ACC neurons (p=0.002, One-Way ANOVA, Fisher’s LSD post-hoc, Fig. 5G), without altering amplitude or decay time. This data suggests a predominantly presynaptic effect of CCH, suppressing neurotransmitter release from excitatory axon terminals in ACC.

**Figure 5.**
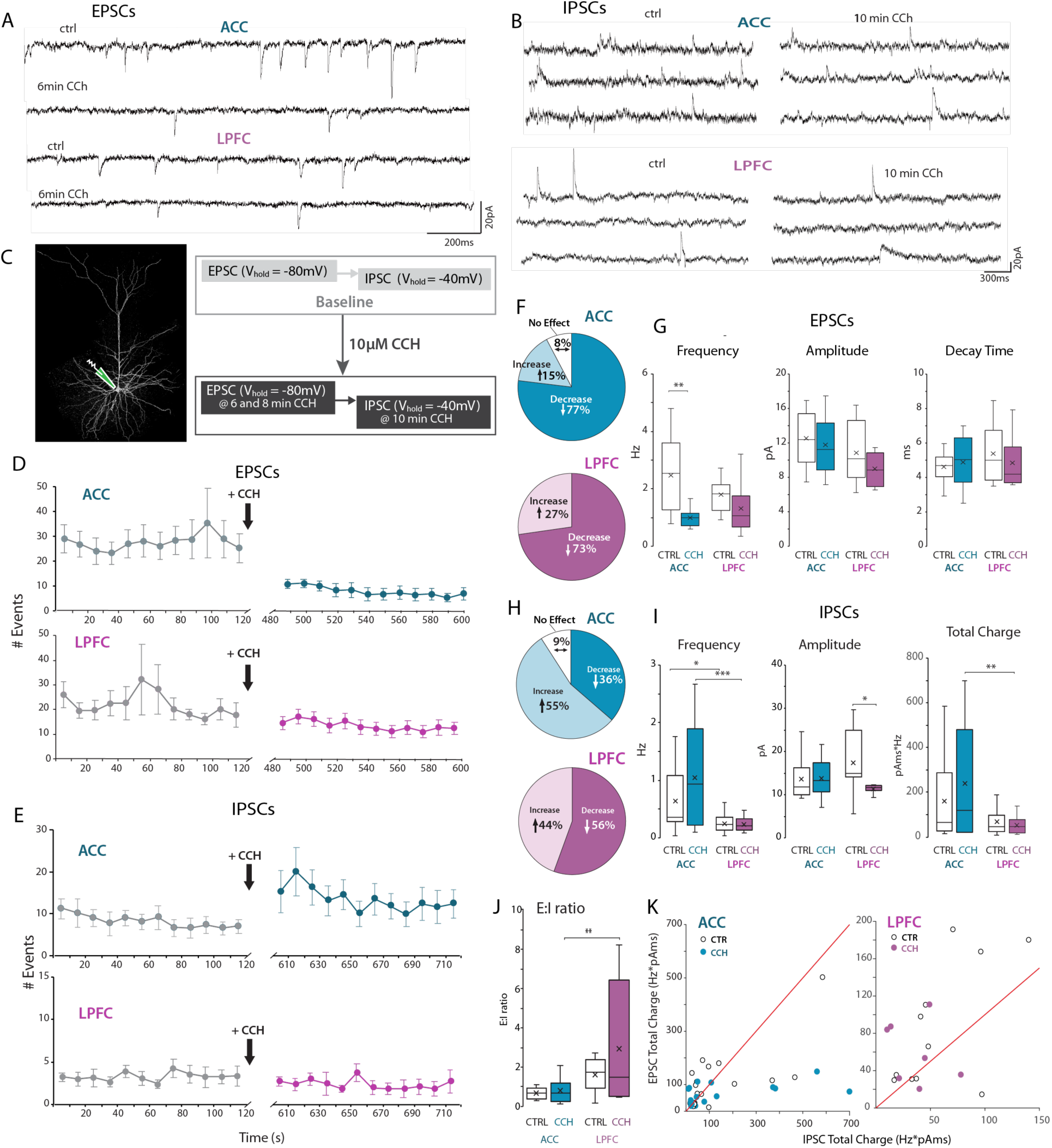
**Cholinergic agonist amplified regional differences in excitatory:inhibitory synaptic tone of ACC vs. LPFC pyramidal neurons**. (**A, B**) Representative traces of (**A**) spontaneous EPSCs (Vhold at-80mV) in ACC (n=12 neurons) and LPFC (n=11); and (**B**) spontaneous IPSCs (Vhold – 40mV) in ACC (n= 11) and LPFC (n= 9) L3 pyramidal neurons, before and after bath application of 10 µM carbachol (CCH), (n=6 cases). (**C**) Schematic of whole cell patch clamp recordings from L3 pyramidal neurons (left) and experimental design for studying synaptic currents (right). (**D, C**) Time-course of number of EPSC and IPSC events (mean per 10 sec time bin) in ACC and LPFC neurons before (control) and during the last 8-10min (D, EPSCs) and 10-12min (E, IPSCs) of bath application of 10µM CCH. Error bars = SEM for each bin. (**F**) The proportion of ACC or LPFC neurons that had a decrease, increase, or no change in the frequency of EPSCs with CCH. (**G**) Box and whisker plots of EPSC properties (frequency, amplitude, decay time) in neurons that exhibited a decrease in EPSC events before and after 8 min in CCH. ** p ≤ 0.01. (**H**) The proportion of ACC and LPFC neurons that had a decrease, increase, or no change in the frequency of IPSCs after CCH. (**I**) Box and whisker plots of IPSC properties [frequency, amplitude, and total charge (frequency x area)] before and after 10 min in CCH. (**J**) The estimated E:I ratio based on the frequency and mean area of each neuron. (**K**) Scatter plots of excitatory versus inhibitory total charge in each ACC (*left*) and LPFC (*right*) neuron from control (open circles) and CCH (filled circles). The red line in both plots represent the linear relationship whereby E=I. In majority of ACC neurons, IPSC total charge dominates, especially after CCH. In LPFC neurons, EPSC total charge dominates. *p ≤ 0.05, ** p ≤ 0.01, *** p ≤ 0.001. Extended Fig. 5 shows washout control experiments.

Analysis of IPSCs revealed heterogeneity in the CCH effect between the two areas (Fig. 5E, H, I). In the ACC, 55% of recorded cells exhibited a CCH-related *increase* in IPSCs frequency, while 36% had a decrease, and 9% exhibited no change from baseline (Fig. 5H). By contrast, in the LPFC, 56% of neurons exhibited a CCH-related *decrease* in IPSC frequency and the remaining 44% displayed an increase. No significant CCH-related differences in mean IPSC frequency were observed in either ACC or LPFC, likely due to the heterogeneity in the CCH effects (Fig. 5E, I).

However, consistent with our previous work ^27^, ACC L3 pyramidal neurons had a significantly higher frequency of IPSCs compared to LPFC, and this difference was enhanced by CCH (p<0.04, Fig.7I). Further, CCH also potentiated the inter-regional difference in total IPSC charge transfer (frequency x area), which was greater in ACC than LPFC neurons (p= 0.008, Fig. 5I). Interestingly, we found that CCH significantly decreased the amplitude of IPSCs only in LPFC pyramidal neurons (control vs CCH, p=0.012), but had no effect on IPSC kinetics (Fig. 5I). These data revealed heterogenous effects of CCH on IPSCs that likely involved distinct inhibitory neurons. Control washout experiments in a subset of EPSC and IPSC recordings demonstrate the reversal of CCH effects with washout to baseline (Extended Fig. 5A, B).

Due to the predominance of inhibition in ACC, we found that the total excitatory to inhibitory charge transfer (E:I) ratio was significantly lower in ACC than in LPFC neurons, especially after CCH exposure (p=0.005, Fig. 5J). Scatter plots of EPSC versus IPSC for ACC and LPFC (Fig. 5K) show that, in ACC neurons, before and after CCH exposure, the E:I ratios were <1, suggesting a predominance of inhibition over excitation; However, in LPFC neurons E:I ratios were >1, thereby suggesting a predominance of excitation over inhibition (Fig. 5J, K). These data show that CCH can differentially shift the E:I balance in ACC and LPFC.

Previous studies in rodents show that cholinergic activation can modulate plasticity of spines – the postsynaptic sites of excitatory synapses^87–89^. We therefore assessed spine density and morphology, which reflect different states of plasticity^80, 82–84, 90^, in L3 pyramidal neurons treated with or without CCH. A two-way ANOVA (treatment group x area) revealed a significant main effect of CCH treatment on subtype distribution and size of spines on apical and basal dendrites of ACC and LPFC neurons (Fig. 6A). In both areas, compared to control, CCH-treated neurons had significantly lower mean densities and proportions of mushroom and stubby spines and a concomitant increase in the proportion of thin spines, on apical and basal dendrites (main effect, treatment, p < 0.001, Fisher’s LSD post-hoc p < 0.05; Fig. 6C). Significant between-area differences were only found on basal dendrites, with densities of total and thin spines greater in ACC compared to LPFC neurons in both treatment groups (main effect, area p < 0.001, post-hoc p < 0.05; Fig. 6B).

**Figure 6.**
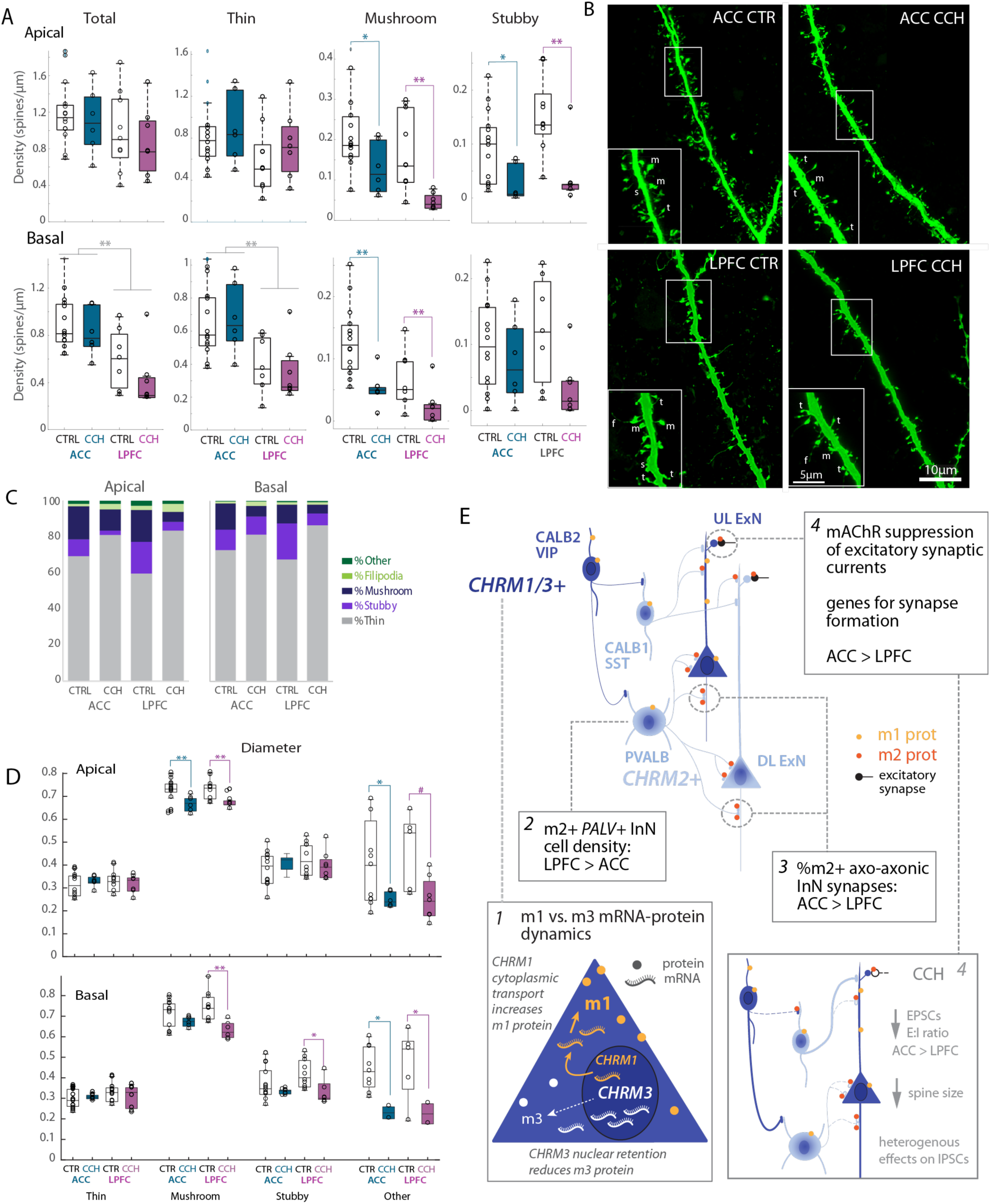
**Cholinergic agonist promoted motile post-synaptic spine morphologies in ACC and LPFC pyramidal neurons**. (**A**) Box and whisker and vertical scatter plots of spine density (total and by subtype) on apical and basal dendrites of individual control and CCH-treated L3 pyramidal neurons in ACC and LPFC. Significant main effect of treatment on apical mushroom (p = 0.0006) and stubby spines (p = 0.0002), and basal mushroom spines (p = 0.001; Fisher post-hoc: ACC: Ap mushroom, stubby, p = 0.03; Bas mushroom p = 0.002; LPFC: Ap mushroom p = 0.004, stubby p = 0.001; Bas mushroom; p = 0.008). Significant main effect of area on basal total (p = 0.0003) and thin spines (p= 0.0004, Fisher LSD post-hoc p < 0.0005). (**B**) Representative z-maximum projections of confocal images of mid-apical dendrites of control and CCH treated ACC and LPFC L3 pyramidal neurons. The neurons were filled with biocytin and labeled with Streptavidin-Alexa 488. (**C**) Average proportion of spines per subtype on apical and basal dendrites of control (open bars) and CCH treated (filled bars) L3 pyramidal neurons in ACC and LPFC, as represented by stacked bar graphs (Fisher’s exact test, p < 0.0001). (**D**) Box and whisker and vertical scatter plots of the average major diameter of spine subtypes on control and CCH treated pyramidal neurons. *p ≤ 0.05, ** p ≤ 0.01, *** p ≤ 0.001. (**E**) Summary schematic of findings showing cell-specific *CHRM1/3*+ vs *CHMR2+* ExNs and InNs enrichment patterns, associated potential differences in mRNA-protein dynamics and trafficking of m1 vs m3 (1), and regional differences in downstream functional effects of ACh on synaptic plasticity and function (2), within the context of known laminar and regional differences in excitatory and inhibitory connectivity, cellular composition (3) and mAChR subcellular localization (4).

Between-treatment differences were also found for spine size, with smaller head diameter of apical mushroom spines in CCH compared to control neurons in both areas (main effect, treatment, p < 0.001; CCH vs CTRL post-hoc: ACC p = 0.01, LPFC p = 0.0005, Fig. 6D, E). In basal dendrites, this CCH-related decrease in spine head size was apparent only for LPFC neurons (mushroom p = 0.002, stubby p = 0.03, Fig. 6E). In addition, spines that were either branched or emerging from other spines (spinules) of CCH treated neurons also exhibited smaller diameters relative to control in both areas (“other” ACC apical/basal p = 0.01; LPFC basal p = 0.03; Fig. 6D, E). Overall, these data suggest that cholinergic stimulation with CCH can lead to shifts in spine morphology and plasticity^80, 82–84, 90^.

## Discussion

### CHRM3 is the most widely expressed mAChR gene in both the ACC and LPFC, despite m1 receptors being predominant at the protein level

Our snRNA-seq analysis of mAChR expression across neuronal subtypes in the macaque frontal cortex revealed intriguing patterns that both align with and diverge from previous protein-level studies^47, 91, 92^. Consistent with earlier work^40, 41^, we found that most cortical neurons co-express m1 and m2 receptors at the protein and mRNA level. However, the gene expression hierarchy (*CHRM*3>*CHRM*2>*CHRM*1) contrasts with the protein-level predominance of m1 receptors in the cortex. This observed mRNA-protein discrepancy highlights the complex relationship between transcription and translation. Recent research has shown that actively transcribed genes may have fewer RNA copies due to instability, while more stable transcripts can accumulate long after transcription has ceased (reviewed in ^93, 94^). Our combined IHC and FISH experiments revealed a higher proportion of cells expressing m1 protein without detectable *CHRM1*, which could indicate dynamic mRNA transport and turnover with a more stable and longer protein half-life^95^ (Fig. 6E-1). These data are consistent with recent work in macaques showing a lack of correlation between *CHMR1* mRNA and autoradiographic measures of M1 receptor density^96^. In line with this, m1 receptor has been shown recently to undergo SUMOylation which increases protein stability and cell surface expression^97^. Conversely, our finding of *CHRM3* presence in cells lacking detectable protein suggest more dynamic regulation and rapid turnover of m3 receptor protein expression^98^. Indeed, specific phosphorylation patterns can promote receptor internalization, leading to either recycling or degradation, thus impacting overall receptor abundance^99^.

Notably, our data revealed that the subcellular localization patterns of *CHRM1* and *CHRM3* transcripts differed. The predominant cytoplasmic localization of *CHRM1* suggests active translation, while the primarily nuclear localization of *CHRM3* implies regulatory mechanisms involving mRNA retention due to the interaction with RNA-binding proteins^100^ (Fig. 6E-1). Nuclear retention of mRNA could diminish protein abundance^101, 102^. This differential localization might explain, at least partially, the discrepancy in the mRNA and protein levels of the receptors and contribute to their distinct signaling cascades in cortical circuits.

### Transcriptional similarity between CHRM1+ and CHRM3+ ExNs aligns with functional overlap of m1 & m3 belonging to the same pharmacological subclass

Our data revealed a high transcriptional similarity between *CHRM1+* and *CHRM3+* ExNs (evidenced by the lack of DEGs), which are distinct from *CHRM2+* ExNs (Fig. 6E). This similarity reflects the fact that m1 and m3 mAChRs are highly co-expressed and belong to the same M1/M3 pharmacological class, suggesting functional overlap^66^. The M1/M3 class elicit post-synaptic depolarizing effects when bound to ACh, via activation of G_q/11_ proteins and opening of voltage-gated K^+^ and Ca^2+^ channels [^66^; reviewed in ^10, 103, 104^]. Indeed, our transcriptional dataset showed *CHRM1/CHRM3*+ ExNs display upregulation in genes encoding muscarinic sensitive voltage-gated “M-current” K^+^ channels (*KCNQ3*), and R-type high voltage activated Ca^2+^ channels (*CACNA1E)*^105^. M1/M3-activated pathways enhance activity and can trigger downstream spine formation and LTP^9, 87, 88^, consistent with the enrichment of functional terms related to axon guidance, adhesion, and synaptic plasticity in *CHRM1+/CHRM3+* ExNs.

In contrast, m2 and m4 receptors belong to the same M2/M4 mAChR pharmacological class that couples to G_i/o_ proteins that inhibit adenyl cyclase, decreasing cAMP and neurotransmitter release^34, 37, 38^ [reviewed in^9, 10, 103, 104^]. Consistent with this, adenyl cyclase *ADCY1* is downregulated in *CHRM2+* as compared to *CHRM1+ and CHRM3+* ExNs. Furthermore, *CHRM2+* ExNs show upregulation in genes encoding proteins that inhibit glutamate release and action. As such, *GIRK1* encodes for inwardly rectifying K+ channel that hyperpolarizes neurons to inhibit activity^106^, and *GRM3/4* encode mGLUR3/4, metabotropic glutamate receptors linked to presynaptic inhibition of glutamate release^107^. Notably, this predisposition for presynaptic suppression is coupled with enrichment of genes related to synapse stability and formation. In both ACC and LPFC, *CHRM2+* ExNs, were enriched in DEGs coding for positive regulators of synapse maturation and stability *(UNC5D, EPHA5/7, SEMA3A, DCC, TRPC3*)^69, 70, 74–77, 80^. Our data highlight the ability of muscarinic receptors to shape functional connectivity by tuning excitability and synaptic transmission.

#### CHRM1/CHRM3 *vs.* CHRM2 enrichment pattern aligns with layer-specific and neurochemically distinct subclasses of ExNs and InNs

The particularly high degree of *CHRM1/CHRM3* expression in UL2-3 ExNs and *VIP+*/*CALB2+* InNs shown here (Fig. 6E) suggest a predominance of ACH-mediated increase in excitability in these subpopulations^88, 89, 108^. In contrast, DL5-6 ExNs and *PVALB+* InNs exhibited a particularly high enrichment of *CHRM2+* cells, which likely traffic m2 receptors down axons to mediate pre-synaptic effects on glutamatergic and GABAergic neurotransmission^34, 37, 38, 40, 108, 109^. These distinct mAChR expression profiles align with layer-specific ExNs and neurochemical InN subclasses with divergent developmental origins and unique synaptic functions (UL vs. DL ExNs; *VIP/CALB2* caudal vs *PVALB* medial ganglionic eminence InNs^55, 110, 111^). Indeed, our functional data showed heterogenous effects of ACh on InN synapses, likely reflecting involvement of these diverse neurochemical InN subclasses^40^. Transcriptomic data suggest that ACh may suppress strong *PVALB*-mediated somatic inhibition and increase firing activity of ‘disinhibitory’ *VIP+*/*CABL2+* (Calretinin) InNs^52, 53, 56^. Interestingly, InNs expressing the parvalbumin protein (PV+) are more numerous in LPFC than ACC, where other InN subtypes predominate^27, 112, 113^ (Fig. 6E-2). This regional diversity in cellular composition of InNs may confer diverse regional cholinergic effects on network dynamics of ACC and LPFC.^28, 114^

### ACC showed upregulation in genes promoting synapse formation and encoding glutamate receptors and Ca2+ channels as compared to LPFC

While our snRNA-seq data revealed relatively similar distribution of *CHRM*-enriched cells in ACC and LPFC, the functional effects of mAChRs largely depend on subcellular localization of proteins, as well as, cholinergic innervation, which are distinct between these regions^12, 13, 40^.

Compared to LPFC, ACC receives denser cholinergic inputs from the basal forebrain^12, 13^ and exhibits greater density of mAChRs on dendrites of pyramidal ExNs^40^. Assessments of the top between-region DEGs in *CHRM1+/CHRM3+* ExNs revealed significant upregulation in ACC compared to LPFC of genes that promote synapse formation^69, 70, 74–77, 80^, and encode for glutamate ionotropic receptor subtypes and Ca channels that potentiate synaptic responses^82–84^. Similar patterns were observed for *CHRM2+* ExNs, however, there was more pronounced enrichment of terms related to LTD and Sodium Ion transport than in *CHRM1/3+* ExNs. This is interesting since ACh is a robust modulator of limbic structures, such as the ACC, playing a key role in arousal and memory^3–6^. Further, a subset of DEGs upregulated in ACC relative to LPFC *CHRM2+* ExNs (*SCN9A*, *PRKG1*) are implicated in pain signaling^85, 86^, a function attributed to the ACC^115, 116^. As these differences between regions may be affected with age, it would be interesting to explore the effects of aging on our dataset in future studies.

Aligning with between-region DEGs related to synaptic function and plasticity, our functional profiling of synaptic properties in L3 pyramidal neurons revealed regional differences in their responses to the cholinergic agonist CCH (Fig. 6E-4). Consistent with our previous work^27^, we found that ACC neurons exhibited lower E:I ratios than LPFC, which was further amplified by CCH. In ACC, CCH significantly reduced EPSC frequency, indicating ACh-mediated presynaptic suppression of excitatory transmission, thereby unmasking and strengthening inhibitory tone. In contrast, CCH caused no significant changes in EPSC properties in LPFC, but specifically reduced the inhibitory current amplitude, shifting the LPFC microcircuit towards a stronger excitatory tone. These data and our previous work at the protein level^40^ support the heterogenous ACh modulation of neurochemically distinct InNs, likely via ACh suppression of *PVALB+* mediated inhibition, while enhancing ‘disinhibitory’ *CALB2/VIP+* effects. Our data showed that majority of cells in ACC exhibited a CCH-related increase, while in LPFC exhibited a decrease in IPSC frequency.

These data align with the higher density of PV+ neurons in LPFC that express m2, compared to the predominance of non-PV+ InNs in ACC^40^ (Fig. 6E-2). However, when assessing PV+ terminals at the protein level specifically, the proportion of PV+ terminals expressing m2 was greater in ACC than in LPFC (Fig. 6E-3), suggesting that CCH-related increase in IPSCs in ACC likely involve non-PV+ InNs^40^. Moreover, these m2+/PV+ terminals are mainly axo-axonic^40^ in ACC, but are mostly peri-somatic targeting in LPFC, further supporting cholinergic modulation distinct InNs in the two areas. The functional impact of greater absolute density of m2+/PV+ InNs in LPFC, but higher proportion of m2+/PV+ axo-axonic terminals in ACC remains unclear and require future computational models and empirical work selectively activating InNs.

On the structural level, cholinergic stimulation via CCH was associated with a decrease in large mushroom spines, suggesting enhanced spine motility^80, 82–84, 90^, in both areas. However, despite stronger CCH-mediated functional presynaptic suppression in ACC, the CCH-mediated structural changes in spines were more widespread in LPFC, occurring on both the apical and basal dendrites of pyramidal neurons. This regional difference is intriguing given our snRNA-seq data showing upregulation in ACC of several genes associated with synapse stability, including Roundabout pathway genes (*SLIT3, SEMA3A*) and adhesion molecules (*EPHA6, CHD13, NLGN1, ERC1*^69, 70, 74–77, 80^. Cholinergic stimulation of spine motility can result from either increased excitability via m1/m3 receptor activation, or as a compensatory response to decreased pre-synaptic transmission via m2 receptors^9, 10, 87, 88, 109^. It is possible that the pronounced cholinergic suppression of presynaptic transmission in ACC does not translate to postsynaptic changes due to the enrichment of synapse stability pathways. Further, CCH also engage nicotinic receptors, potentially counteracting mAChR-mediated suppression in ACC^117^. Presynaptic alpha7-containing nicotinic receptors can enhance glutamatergic release^118, 119^, and likely contribute to the heterogenous effects of CCH in ACC and LPFC. Receptor-specific pharmacological tools will be crucial to clarify the distinct contributions of ACh receptors to region-specific synaptic efficacy and plasticity.

### Implications for network dynamics and behavior

Our data revealed that mAChR gene enrichment in ACC and LPFC are associated with between-region transcriptional differences in synapse plasticity pathways that aligned with functional experiments. The relative upregulation of genes promoting synapse formation in *CHRM+* ExNs and more pronounced cholinergic suppression of excitatory synaptic currents in ACC, suggests a role of ACh for filtering and enhancing signal-to-noise (gain) of inputs, which can be at play during ACC-mediated tasks requiring high cognitive demand^7, 24, 30–32, 120, 121^. Concomitantly, cholinergic increase in E:I ratio in LPFC may support working memory representations^122, 123^. The role of ACh in gain modulation has been observed in macaque primary visual cortex^124^, rodent prefrontal and olfactory corticesand medial temporal lobe areas [reviewed in^9–11, 25^]. Our findings support the role of ACh as a modulator of synapse plasticity and E:I circuit dynamics within ACC and LPFC, which can endow flexibility for processing of signals depending on the cholinergic tone and cell-specific activation of distinct muscarinic receptors.

## Conflict of Interest Statement

All authors declare that they have no conflicts of interest.

## Acknowledgments

We thank Drs. Douglas Rosene and Tara Moore for access to resources and providing tissue for the experiments; Bethany Bowley, Samantha Calderazzo, Alexander Hsu, Eli Shobin, Penny Shultz Ajay Uprety, Katelyn Batterman, Karen Bottenfield, Karen Slater, and Veronica Go for technical assistance during perfusion and brain cutting. We also thank Dr. Yuriy Alekseyev and the members of the Boston University Chobanian & Avedisian School of Medicine sequencing core for their help and guidance with the sequencing experiments We are grateful to NIH agencies for their support: NIH/NIMH R21MH126250 (MPI: M. Medalla and E. Zeldich), NIH/NIA R21AG072069 (MPIs: J. I. Luebke and E Zeldich), NIH/NIMH R01 MH116008 (PI: M. Medalla); NIH/NIA: R01-AG059028 (MPIs: J. I. Luebke and P. Hof), R01LM013154 (PI: J.D.C), and RF1-AG043640 (PI: D. L. Rosene).

## Author contributions

AT, CAM designed and performed aspects of all the experiments and aided in the bioinformatics analysis, interpreted the results, and wrote the first drafts of the paper. MM, EZ, and JL conceived and oversaw the project, designed and performed aspects of all experiments and analyses, provided guidance, interpreted the results, and wrote the paper. SDK, RY, SA, and JDC designed and performed bioinformatics and statistical analyses for snRNA-seq dataset and edited the paper. BJS, WC, TGV, JG, AC, ILT, JM helped perform experiments, gather and analyze data, and edited the manuscript.

## Data availability

Raw and processed scRNA-seq data generated in this study have been deposited in the NCBI Gene Expression Omnibus database (GEO). Source data are provided in the Source Data files in Supplementary Materials.

## Code availability

The R scripts used for data analysis are available on GitHub [https://github.com/campbio-manuscripts/Muscarinic_snRNAseq_ePhys].

## Competing interests

The authors declare no competing interests.

## Materials & Methods

### Experimental subjects

Brain tissues used in this study were from 10 adult rhesus monkeys (*Macaca mulatta,* n=4 males, ages = 8.5-24.8 y for snRNAseq; n = 4 males and n = 2 females, ages = 8.5-16.8 y for electrophysiology) that were subjects in larger studies of brain aging and cognition (Supplementary Table S1). Monkeys were obtained from either National Primate Centers or private vendors and were individually housed in the Laboratory Animal Science Center (LASC) at Chobanian and Avedisian Boston University School of Medicine (BUSOM). The LASC is fully accredited by the Association for Assessment and Accreditation of Laboratory Animal Care, with animal research conducted in strict accordance with guidelines of the National Institutes of Health’s *Guide for the Care and Use of Laboratory Animals* and the *U.S. Public Health Service Policy on Humane Care* and the Boston University Institutional Animal Care and Use Committee.

### Tissue collection for acute slice preparation and molecular experiments

Tissue was harvested using a two-stage transcardial perfusion protocol that allowed for the harvest of live tissue prior to completing the perfusion fixation of the remaining brain tissue ^125^. The animals were initially sedated with ketamine hydrochloride (10 mg/kg) and deeply anesthetized with sodium pentobarbital (to effect, 15 mg/kg, i.v), followed by a two-stage transcardial perfusion that begins with ice-cold (4°C) Krebs–Henseleit buffer to clear the vasculature and slow proteolysis to allow for the harvest of fresh tissue from the left hemisphere for molecular and electrophysiological experiments. The first block of tissue (∼10 mm^3^) was removed first from the LPFC, the caudal half of the ventral bank of the principal sulcus equivalent to area 46 ^126^. The second block of the same size was removed from the ACC, within the rostral part of A24 in ventral bank of the cingulate sulcus ^126, 127^. Each block was transferred into oxygenated (95% O_2_, 5% CO_2_) ice-cold Ringer’s solution (in mM: 26 NaHCO_3_, 124 NaCl, 2 KCl, 3 KH_2_PO_4_, 10 glucose, and 1.3 MgCl_2_, pH 7.4), and tissue was vibratome-sectioned for electrophysiological recordings or flash frozen in liquid nitrogen and stored at-80°C for single nucleus RNA sequencing experiments. Upon completion of the tissue harvest, the perfusate was switched to 4% paraformaldehyde in 0.1M phosphate buffer (PB, pH 7.4, 37°C) to fix the remaining brain tissue. The brain was then carefully removed, weighed, and post-fixed in 4% paraformaldehyde prior to being cryoprotected through a series of glycerol solution. The brains were ultimately flash-frozen in-70°C isopentane (Rosene et al., 1986) and stored at-80°C until they are cut on a freezing microtome in the coronal plane at 30 and/or 60µm sections and stored in 15% glycerol buffer for tissue analysis.

### Processing brain tissue samples for single nucleus RNA sequencing (snRNA-seq)

ACC and LPFC tissue samples that were previously flash-frozen and stored at-80°C were dissociated by Dounce homogenization in lysis buffer (250mM sucrose, 25mM KCl, 5mM MgCl_2_, 10mM Tris buffer, pH 8.0, 0.1% Triton X-100, 1µM DTT, 1x Protease inhibitor, 1x RNase inhibitor and 1µM DAPI). The homogenate was then filtered through a 40µm-cell strainer to collect single nuclei. Fluorescence-activated cell-sorting (FACS) was performed at the Chobanian and Avedisian BUSOM Flow Cytometry Core Facility on the BD FACS ARIA II SORP (Becton Dickson) to remove cellular debris and to capture singlets by gating on forward and side-scatter light properties and nuclei fluorescent expression of DAPI. We collected 50,000 nuclei into a collection buffer (0.04% Bovine Serum Albumin (BSA), in PBS supplemented with 1x Protease inhibitor, and 1x RNase inhibitor) as previously described ^128^.

The sorted nuclei were then processed at the Chobanian and Avedisian BUSOM Microarray and Sequencing Resource Core Facility using the Chromium droplet-based platform (10X Genomics) whereby single nuclei, reagents, and a single gel bead containing barcoded oligonucleotides are encapsulated into nanoliter-sized Gel Bead-in-Emulsion (GEMs) using the GemCode platform for downstream reverse transcription of RNAs. Full length, barcoded cDNA was then amplified by polymerase chain reaction (PCR) for the generation of snRNA-seq libraries. The resulting cDNA libraries were assessed via Bioanalyzer High Sensitivity DNA Assay (Agilent Technologies, USA) and sequences on an Illumina NextSeq500 instrument with the target capture of 2000 cells per sample in accordance with the Illumina and 10x Genomics guidelines with 1.4-18.8 input and 1% PhiX control library spike-in (Illumina, USA).

### snRNA-seq processing and analyses

The sequencing reads were processed with the 10X Genomics Cell Ranger V3 pipeline^129^ using a modified reference that included both introns and exons regions to generate a unique molecular identifier (UMI)/feature-barcode matrix. Quality control was performed using Seurat^130^ and singleCellTK^131^ packages. Nuclei were excluded based on lower numbers of UMIs detected (≤200) or high percentage of counts derived from mitochondrial genes (≥7.5%). Features that were detected in fewer than 3 cells were excluded.

The Seurat package was also used for clustering while normalization was performed with the NormalizeData function. Variable features were selected with the FindVariableFeatures function using the “vst” method. Principal component analysis (PCA) was performed using the top 2,000 most variable genes with the RunPCA function. Clustering of cells was implemented with the FindClusters function using 35 PCs. Cell types were annotated based on canonical markers (Supplementary Table S2).

The enrichment of RNA expression of markers of interest (*CHRM*1 *CHRM2*, *CHRM*3, *CHRM4*) among the cell clusters in the ACC and LPFC was calculated based of the proportion of cells expressing these genes. Differential gene expression analysis was performed on the clusters of excitatory (ExN) and inhibitory (InN) neurons to assess differentially expressed genes (DEGs) between ACC and LPFC, as well as among subpopulations of neurons enriched for muscarinic receptors, using Seurat’s FindMarkers function. Lastly, using the list of DEGs from each comparison, we performed a gene-ontology (GO) analysis using a gene set search engine Enrichr (https://maayanlab.cloud/Enrichr/)^61^. Post-analysis filtering was used to cutoff at an adjusted p-value of p<0.05 and a false discovery rate (FDR) less than 0.25.

### Immunohistochemistry and HCR-FISH

To validate our snRNA-seq data, we performed combined immunohistochemical (IHC) and fluorescent in situ hybridization using hybridization chain reaction (HCR)^132, 133^ on serial coronal sections of archived tissue from 4 monkeys. Prior to IHC-FISH experiments, frozen 30µm tissue sections were post-fixed in 4% PFA for 2 hrs at RT. Sections were then rinsed (3 x 10 min, RT) in 0.01M RNase-free phosphate-buffered saline (PBS; Gibco) and stored in filtered 70% RNase-free, molecular grade ethanol in 4°C until use. IHC was performed to visualize and quantify the expression of m1 and m3 cholinergic receptor proteins, as previously described^40, 41^. Briefly, sections were incubated in 50mM glycine in 0.01 M PBS, pH 7.4, to quench free aldehydes from the fixation step. The sections were then rinsed in 0.01 M PBS (3 x 10 min, RT) and incubated in 10mM sodium citrate, pH 8.5, in a 70°C water bath for 20 min for antigen retrieval. After rinsing in 0.01 M PBS (3 x 10 min, RT), the sections were incubated in a blocking solution [0.01M PBS, 5% bovine serum albumin (BSA), 5% normal donkey serum (NDS), 0.2% Triton X-100] to reduce non-specific binding of secondary antibodies. Sections were then incubated with gentle agitation at 4°C for 48 hrs in a solution with primary antibodies against MAP2, m1, and m3 receptors [diluted in 0.1M PB, 0.2% acetylated BSA (BSA-c, Aurion), 1 % NDS, 0.1 % Triton X-100; Supplementary Table S3]. To enhance antibody penetration, sections were microwaved for two sessions (2 x 10 min, 150W, 37°C per session) in low-wattage PELCO Biowave (Ted Pella Inc.). After incubation with primary antibodies, the sections were rinsed in 0.01 M PBS (3 x 10 min, RT), were microwaved for a single session (2 x 10 min, 150W, 37°C), and were incubated with gentle agitation at 4°C overnight in a solution with secondary antibodies (Supplementary Table S3). Prior to HCR-FISH, sections were rinsed (3 x 10 min, RT) in 0.01M PBS, post-fixed with 4% PFA for 30 min at RT, and dehydrated with 70% ethanol for 30 min at 4°C. Afterward, sections were rinsed in 2x Ultrapure saline sodium citrate (SSC, Thermo Fisher Scientific) buffer (3 x 1 hr, RT). Sections were then equilibrated with Hybridization Buffer (Molecular Instruments) for 30 min at 37°C before overnight hybridization with HCR Probes for *CHRM1* and *CHRM3* (final concentration of 4nM, Supplementary Table S3) in Hybridization Buffer at 37°C. The next day, sections were rinsed with Probe Wash Buffer (3 x 10 min and 1 x 1 hr, 37°C) and with 5x SSC with Tween-20 (SSCT) (3 x 5 min, RT). Afterward, the sections were equilibrated with Amplification Buffer (Molecular Instruments) for 30 min at 37°C. During the equilibration step, fluorescently labeled HCR hairpins were denatured at 95°C for 90 sec and then cooled to RT for 30 min. The denatured hairpins (final concentration of 60nM) were combined into a new tube with Amplification Buffer and applied to tissue sections for 16 hrs in the dark at RT. The following day, the sections were rinsed with 5x SSCT (3 x 5 min and 1 x 30 min, 4°C) before an overnight incubation at 4°C with fresh 5x SSCT. Sections were then mounted onto Superfrost Plus glass slides (Fisher Scientific), coverslipped with ProLong Gold Antifade Mountant (Thermo Fisher Scientific), and cured at RT in the dark.

### Cell density estimates of m1 and m3 protein and mRNA expressing neurons

As described in our previous work^41^, we quantified the density of somata labeled for m1 and m3 protein and mRNA. Confocal image stacks from fields in L2-3 and L5-6 of ACC area 24 and LPFC ventral A46 (n = 2 fields per layer/area/case) were obtained using laser-scanning confocal microscope (Olympus FV3000). Image stacks were acquired using a 40x/1.4 NA oil immersion objective (Olympus UPlanXAPo) at a voxel resolution of 0.124µm x 0.124µm x 0.5µm. For each area, we identified 1-2 columns per section, and systematically imaged ROIs within L2-3 and L5-L6. We focused on imaging these layers where somata of pyramidal neurons mainly reside, since our goal is to validate expression patterns in ExNs in the snRNA-seq dataset (Barbas and Pandya, 1989). The resulting images were deconvolved to improve the signal-to-noise ratio and converted to 8-bit image files for analysis using cellSens software (Olympus). For each deconvolved image stack, we used semi-automated classifier tools in QuPath ^134^ to quantify the distribution of m1 and m3 cells co-expressing protein and mRNA as described^40^. For each field, the maximum z-projection of three 2 µm substacks of top, middle, and bottom optical slices were generated in Fiji/ImageJ (https://imagej.net/Fiji; RRID:SCR_002285) and imported into QuPath. Cell detection function was used to automatically segment somata of m1 protein^+^ cells as individual objects. The segmented objects were checked for positive expression of neuronal MAP2 to include neurons (and exclude glia). Majority of the strongly labeled MAP2 neurons are excitatory, but lightly labeled non-pyramidal neurons were also included and are likely InN. The segmented objects were then classified based on their expression of m1 mRNA, m3 mRNA, and m3 protein using the semi-automated classifier tools in QuPath. The classifiers were based on a user-defined intensity threshold and was kept consistent within each case. The proportions of m1 and m3 mRNA and protein-expressing neurons were calculated by dividing each classified marker by the total number of detections (m1 protein^+^ or m3 mRNA^+^ and protein^+^ cells).

### Functional assessments of muscarinic receptor activation on spontaneous excitatory (EPSCs) *and inhibitory (IPSCs) postsynaptic currents in layer 3 pyramidal neurons*

In parallel to tissue preparation for biochemical experiments, a part of the fresh tissue blocks harvested from the ACC and LPFC were sectioned into 300µM coronal slices in ice cold Ringer’s solution using a vibrating microtome (Leica VT1000S), as described^27^. The resulting slices were then placed into room temperature, oxygenated Ringer’s and allowed to equilibrate for 1 hour. Following the equilibration periods, individual slices were placed into a submersion-type recording chamber (Harvard Apparatus, Holliston, MA) and mounted onto the stages of Nikon E600 infrared-differential interference contrast (IR-DIC) microscopes (Micro Video Instruments). All experiments took place at room temperature in oxygenated Ringer’s solution (at a rate of 2-2.5mL/min), which improve the viability and duration of recordings from monkey cortical slices.

Neurons were visualized under the IR-DIC contrast optics and electrodes were fabricated on horizontal Flaming and Brown micropipette puller (Model P-87, Sutter Instruments, Novato, CA, USA) and were filled with either potassium methanesulfonate-based internal solution (in mM: 122 KCH_3_SO_3_, 2 MgCl_2_, 5 EGTA, 10 NaHEPES, with 1% biocytin, pH 7.4) with resistances of 3-6 MΩ in the external Ringer’s solution. Data were acquired using EPC-9 or EPC-10 patch-clamp amplifiers using PatchMaster software (HEKA Elektronik, Lambrecht, Germany). Neurons were included in electrophysiological analysis if they had a resting membrane potential of ≤-55mV, stable access resistance, an action potential (AP) overshoot, and repetitive firing responses^125^. Spontaneous excitatory and inhibitory synaptic currents (EPSCs and IPSCs) were recorded for at least 2 minutes at a holding potential of-80mV and-40mV, respectively^27^ followed by a continuous bath application of a non-specific ACh agonist carbachol (CCH, 10µM) and recording of EPSCs and IPSCs for ∼8 minutes total. For some neurons, a 15-minute washout in ringers with continued collection of EPSCs and IPSC was performed after CCH exposure. Analysis of synaptic events was performed using MiniAnalysis (Synaptosoft, Decatur, GA, USA), with the event detection threshold set at maximum root mean squared noise level (5 pA) as described previously.

We examined the frequency, mean amplitude, area under the curve and kinetics (rise and decay time, and half-width).

### Processing of filled neurons for assessments of spine density

During whole-cell patch-clamp recordings, neurons were filled with biocytin (0.5%, Millipore Sigma) and processed for post-hoc morphological assessments, as described in our previous work^27, 135^. To visualize recorded cells filled with biocytin, 300-µm slices were fixed in 4% paraformaldehyde in 0.1 M phosphate-buffered saline (PBS; pH = 7.4), incubated for 2 h in 1% Triton X (in 0.1 M PBS), followed by 48 h in streptavidin-Alexa 488 (diluted 1:500 in 0.1 M PBS; Invitrogen). Slices were washed in 0.1M PB, mounted on glass slides, and coverslipped with Prolong Gold mounting media (ThermoFisher).

### Confocal imaging and spine analyses of filled neurons

Neurons with complete somata and apical dendrites (with no cut branches in the proximal third) were included in morphological analyses. Image stacks were acquired using an Olympus FV3000 confocal laser-scanning microscope (Evident/Olympus Inc). For assessment of filled spines, a series of image stacks were acquired (at 0.04 x 0.04 x 0.3µm per voxel, using 60x/1.5 NA oil-immersion objective, UPlanXApo, Evident/Olympus Inc) with a 488 nm excitation laser. For each neuron, confocal stacks within 100 µm from the tissue surface were acquired to image one complete basilar dendritic branch, and the main apical trunk followed to the end of one complete distal apical dendritic branch. Image stacks are then deconvolved using AutoQuant (Media Cybernetics, Bethesda, MD, USA), and tiled and analyzed using Neurolucida 360 (MBF Bioscience; RRID:nif-0000-10294)^27^.

Subsampled basal and apical dendrites were manually traced and spines along these dendrites were manually marked and classified by subtypes and size, as described previously ^135^.

Spines were classified based on head width and neck length as follows: thin (width >0.6 µm), mushroom (width ≥ 0.6 µm), stubby (lacking a neck), filopodia (neck length ≥ 0.3µm), and other (branched spines or those that cannot be identified). The densities (number of spines per micron of dendritic length) and proportion of spines by subtype were calculated. The measured spine head widths and neck lengths for each cell were also assessed. Sholl analyses with concentric spheres placed at 20-μm increments from the center of the soma ^136^ were performed to determine dendritic and spine parameters of apical and basal arbors as a function of distance from the center of the soma.

---

**Extended Data Figure 1:**
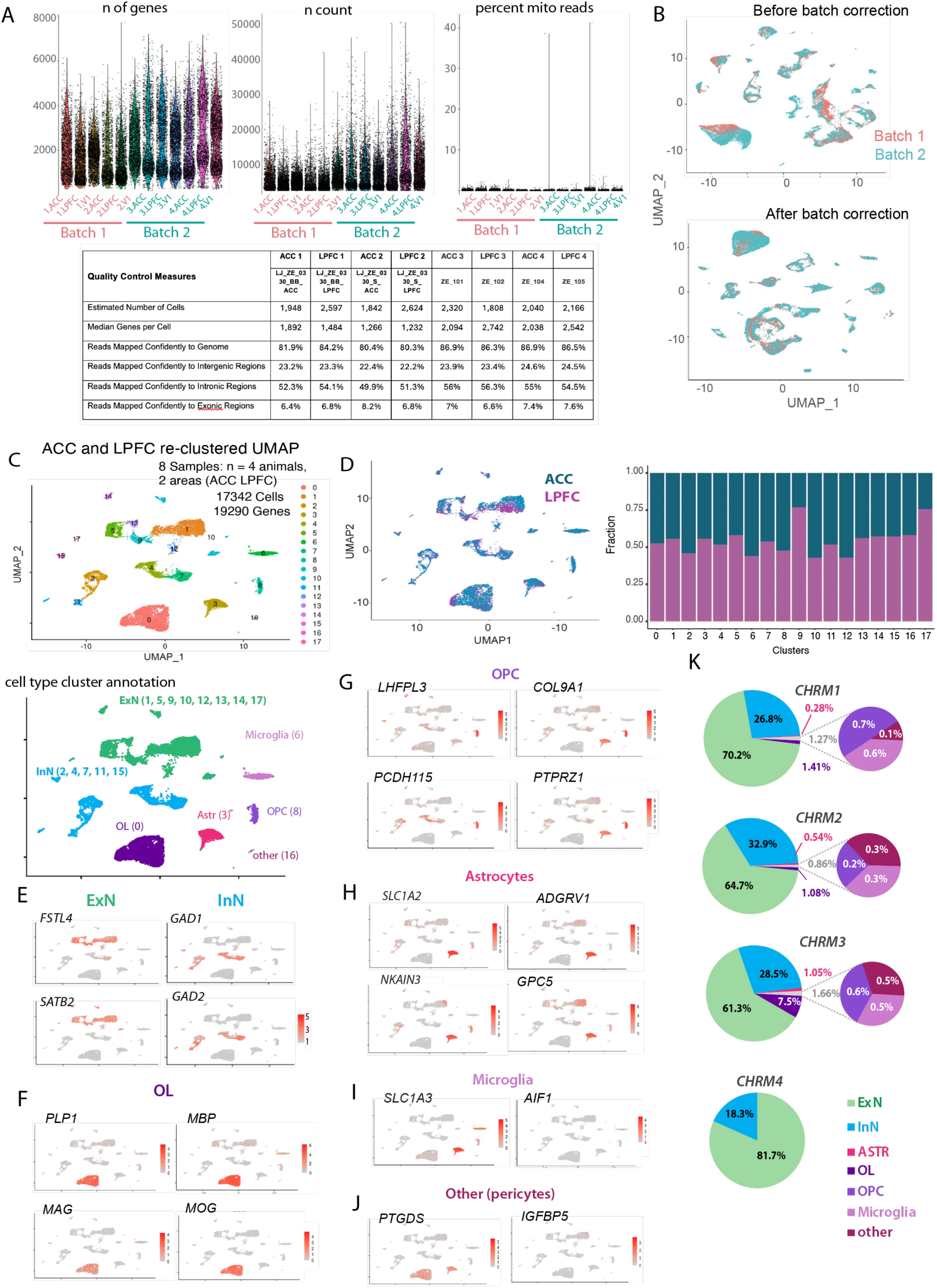
Quality Control, Batch Correction and Annotation Pipeline for snRNAseq dataset. (A) Quality Control: (Top) Scatter plot showing the number of genes (nFeatures), number of RNA molecules (nCount), and the percentage of mitochondrial genes before filtering in each sample collected from the three cortical areas ACC and LPFC with primary visual cortex (V1). (Bottom) Table of quality control assessment of single nucleus RNA sequencing data. Low quality cells with number of genes (nFeature) below 500, cells with number of RNA molecules (nCount) below 800, and with mitochondrial genes above 10% were filtered and exclude from the dataset. After filtering we had 19605 genes and 26650 cells from three brain areas (ACC, LPFC, V1), n = 4 subjects, processed in 2 batches. (B) UMAP plot of nuclei from ACC, LPFC, and V1, through two batches, annotated by batch, to show clustering before and after batch correction. (C) UMAP plot of nuclei from ACC and LPFC re-clustered for further analyses of 19290 genes and 17342 cells from 2 areas, n = 4 monkeys. Bottom shows UMAP of ACC and LPFC clusters annotated by major cell type based on canonical marker expression. (D) UMAP plot of nuclei annotated by area (left), and cell abundance bar plot (right) showing the proportion of cells in each area per cluster. All clusters contained the representation of cell from both regions. (E-K) UMAP feature plots showing expression pattern of a subset of canonical markers used to identify the major cell classes (See Supplemental Table S1): (E) Excitatory (ExN) and Inhibitory (InN) Neurons, (F) Oligodendrocytes (OL); (G) Oligodendrocyte Precursor Cells (OPC); (H) Astrocytes; (I) Microglia and (J) Other cells mainly pericytes. (K) Pie charts showing the distribution of neuronal and glial cell types within the subpopulation of cells expressing each *CHRM* gene from both areas pooled together.

**Extended Data Figure 2:**
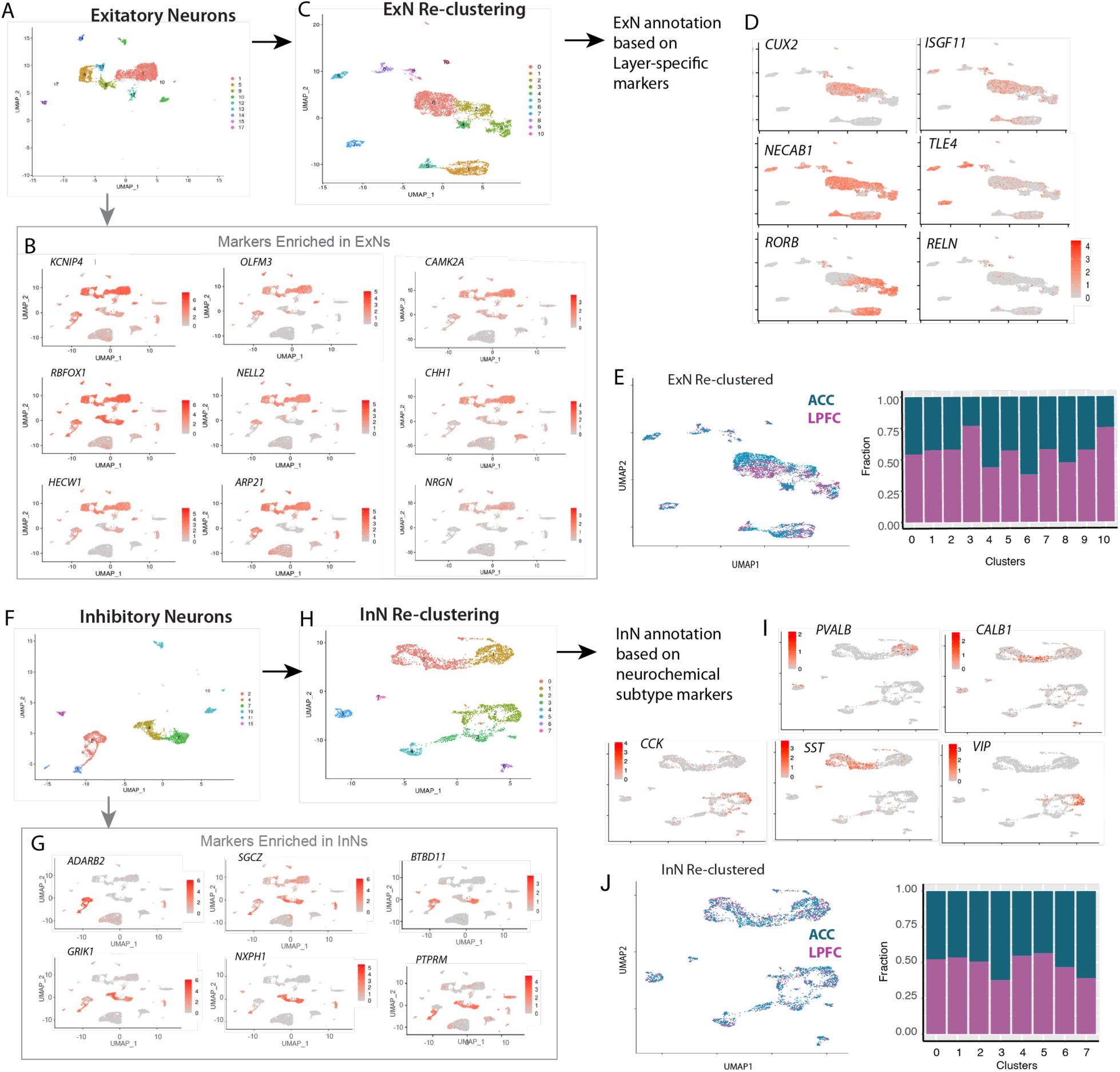
Re-clustering and annotation of ExN and InN subclasses in ACC and LPFC. (A) UMAP plot of the subset of ExN clusters from the full dataset for all cell types. (B) UMAP feature plot of the canonical markers enriched within major cell class annotated as ExNs. (C) UMAP plot of ExNs after re-clustering, annotated by cluster number. (D) UMAP feature plots showing expression pattern of a subset of layer specific markers used to annotate re-clustered ExN by laminar subclass; (E) UMAP plot of ExNs after re-clustering, annotated by cortical region (left) and cell abundance stacked bar plot (right) showing proportion of nuclei from ACC vs LPFC in each ExN cluster. Each cluster contains a representation of both areas. (F) UMAP of the InN clusters from the full dataset for all cell types. (G) Canonical markers enriched within major cell class annotated as InNs. (H) UMAP plot of InNs after re-clustering, annotated by cluster number. (I) UMAP feature plots showing the expression pattern of a subset of specific markers representing functionally distinct neurochemical subclasses based on the literature, used to annotate re-clustered InN by subclass. (J) UMAP plot of InN after re-clustering annotated by cortical region (left) and cell abundance stacked bar plot (right) showing the proportion of nuclei from ACC vs LPFC in each InN cluster. Each cluster contains a representation of both areas.

**Extended Data Figure 3:**
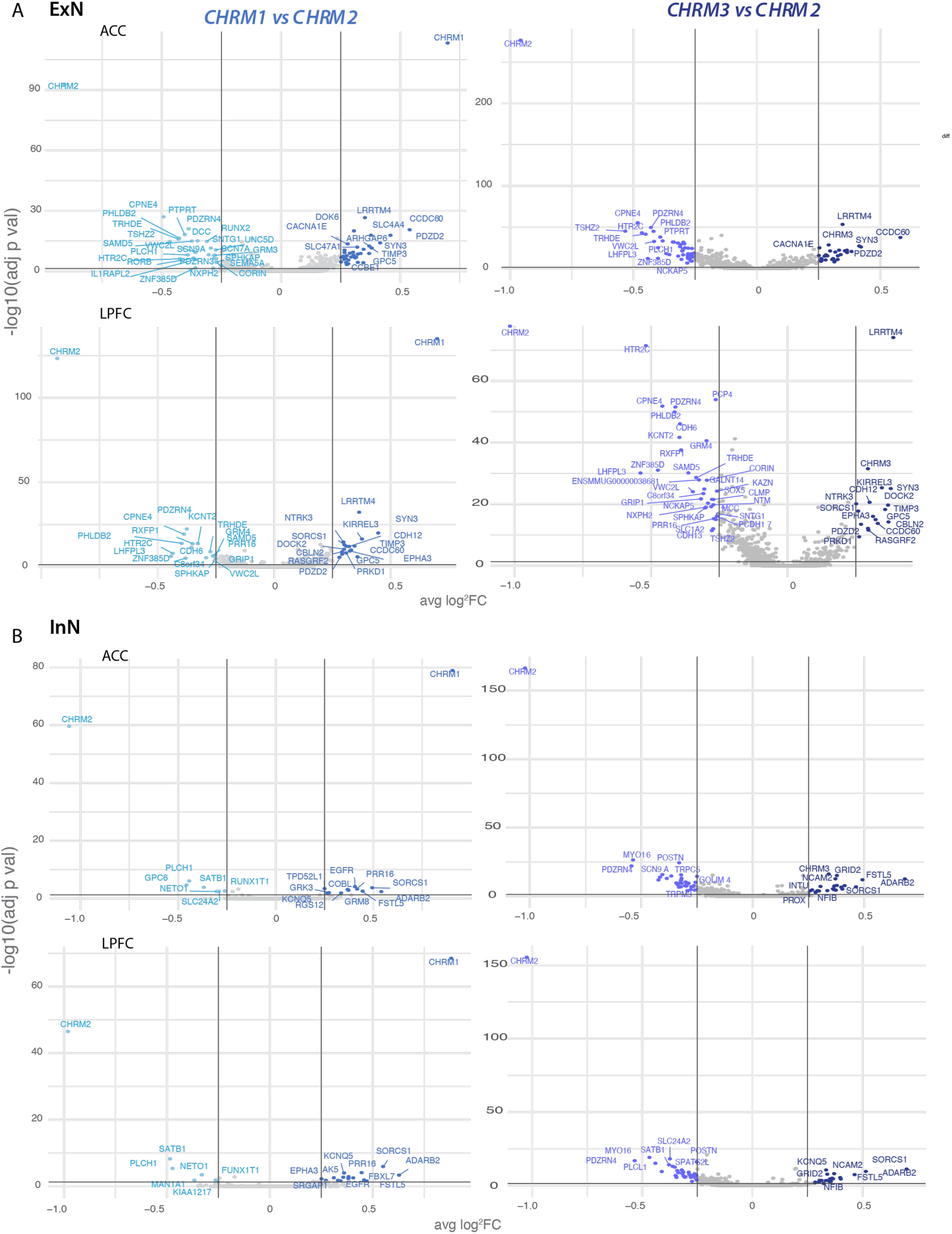
Differentially express genes across *CHRM* expressing neurons in ACC and LPFC. (A) Volcano plots showing DEGs between *CHRM1+* vs. *CHRM2+* and *CHRM3+* vs. *CHRM2+* ExNs in ACC and LPFC. (B) Volcano plots showing DEGs between InNs in ACC and LPFC.

**Extended Data Figure 5:**
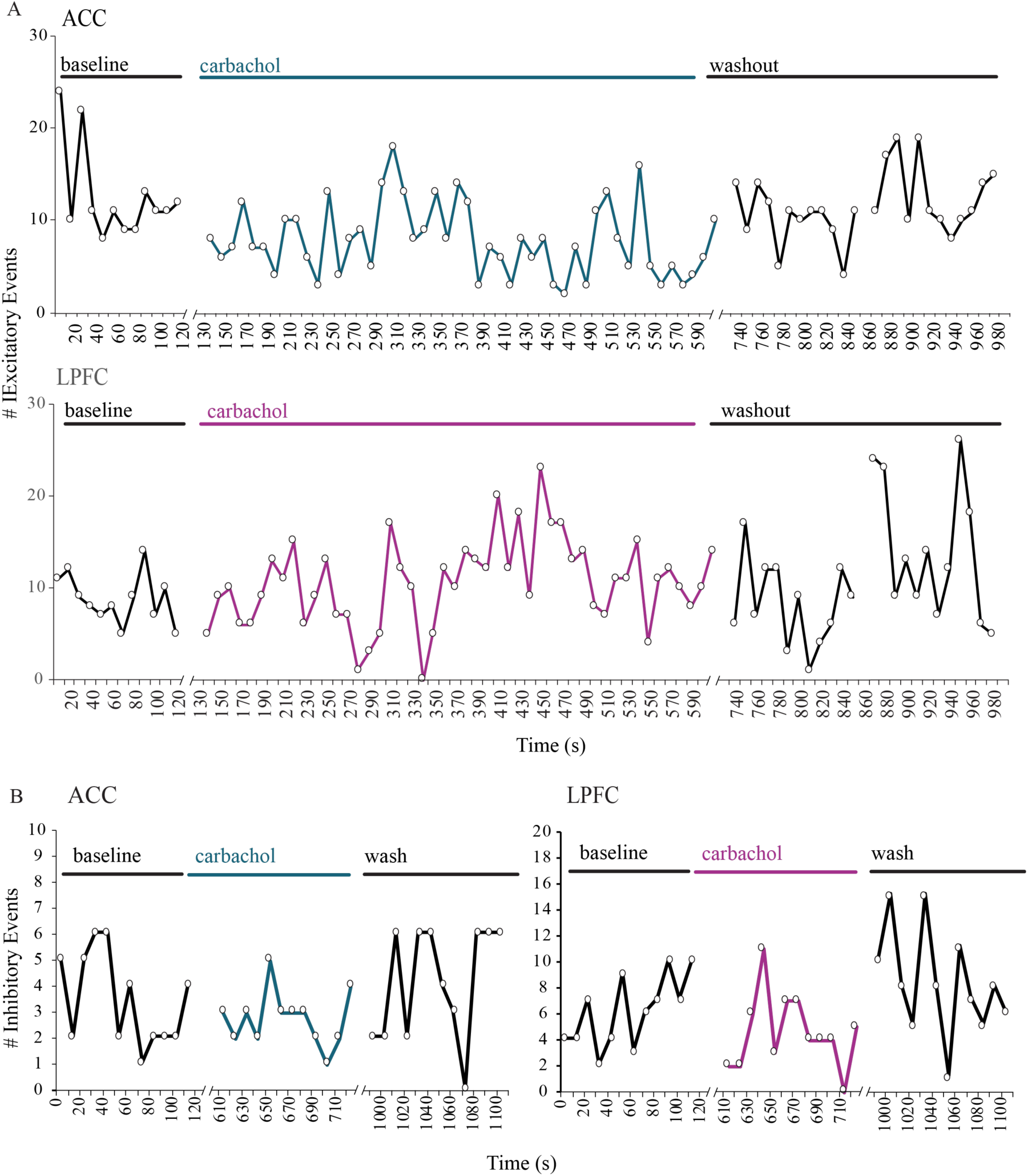
CCH washout experiments during recording of synaptic currents. (A, B) Time course of excitatory (A) and inhibitory (B) synaptic currents in ACC and LPFC to examine the effect before (baseline), after the addition of CCH (carbachol), and during the first 2-minutes of washout and after 15-minute washout period (washout).

## Notes

https://github.com/campbio-manuscripts/Muscarinic_snRNAseq_ePhys

### Competing Interest Statement

The authors have declared no competing interest.

