## Supplementary Tables S1 S2 S3 for "Transcriptional and functional profiles of muscarinic receptor-expressing neurons in primate lateral prefrontal and anterior cingulate cortices"

**Supplementary Data Tables:**

**Table S1. Subjects & Experiments.** Age and sex of rhesus monkeys used in electrophysiology, snRNA sequencing and IHC/HCR-fISH experiments

| **ID Name** | **Sex** | **Age** | **Experiment** |
| --- | --- | --- | --- |
| AM367 | M | 8.5 | Single nucleus RNA sequencing, IHC/HCR-FISH |
| AM368 | M | 8.9 | Single nucleus RNA sequencing, IHC/HCR-FISH, Electrophysiology |
| AM365 | M | 24.8 | Single nucleus RNA sequencing |
| AM372 | M | 20.8 | Single nucleus RNA sequencing |
| AM299 | F | 8.6 | IHC/HCR-FISH, Electrophysiology |
| AM358 | F | 8.4 | IHC/HCR-FISH |
| PIK | M | 8.2 | Electrophysiology |
| AM378 | M | 16.8 | Electrophysiology |
| AM321c | M | 13.4 | Electrophysiology |
| SM036 | M | 10 | Electrophysiology |

**Table S2. Cluster Annotation based on Canonical Markers for Major Cell types and ExN Layer-specific genes**

| **UMAP cluster/s** | **Cell Type** | **Markers** | **References** |
| --- | --- | --- | --- |
| 0 | Oligodendrocytes | *PLP1, MBP, MOG, MAG, RFN220* | (Zhu et al. 2018; Chamling et al. 2021) ^42, 43^ |
| 8 | OPCs | *PTPRZ1, LHFPL3, COLA91, PCDH15* | (Zhu *et al.* 2018; Chamling *et al.* 2021) ^42, 43^ |
| 1, 4, 5 6, 7, 10 | Excitatory Neurons | *CUX2, NECTIN3, ISGF11, RORB, FOXP2, FSTL4, SATB2, KCNIP4, HECW1, FSTL4, NELL2, NRGN, CAMK2A,* | (Zhu *et al.* 2018; Li et al. 2022; Suresh et al. 2023) ^43-45^ |
| 2, 4, 7, 11, 15 | Inhibitory Neurons | *LHX6. GAD1, GAD2,* | (Zhu *et al.* 2018; Suresh *et al.* 2023) ^43, 45^ |
| 3, | Astrocyte | *SLC1A2, SLC1A3, NKAIN2, ADGRV1, GPC5* | (Zhu *et al.* 2018; Suresh *et al.* 2023) ^43, 45^ |
| 6 | Microglia | *AIF1, SLC1A3* | **(Olah et al. 2020) ^46^** |
| 16 | Other (Pericytes) | *SLC13A4, TRPM3, PTGDS, IGFBP5, MFAP4* | (Zhu *et al.* 2018; Suresh *et al.* 2023) ^43, 45^ |
| **Genes associated with laminar subclasses of excitatory neurons** | | | |
| ***Marker*** | **Laminar Designation** | **References** | |
| *RELN* | Upper Layers (L1) | (Zeng H et al. 2012) ^49^ | |
| *CUX2* | Upper Layers (L2, L3) | (Zeng H *et al.* 2012; Lake BB et al. 2016) ^48, 49^ | |
| *RASGRF2* | Upper Layers (L2, L3) | (Zeng H *et al.* 2012) ^49^ | |
| *NECTIN3* | Upper Layers (L2, L3) | (Tomorsky J et al. 2020) ^50^ | |
| *IGSF11* | Upper Layers (L3) | (Zeng H *et al.* 2012) ^49^ | |
| *PCP4* | Deep Layer (L5) | (Zeng H *et al.* 2012) ^49^ | |
| *TLE4* | Deep Layer (L5) | (Zeng H *et al.* 2012) ^49^ | |
| *BCL11B* | Deep Layer (L5, L6) | (Maynard KR et al. 2021) ^51^ | |
| *FEZF2* | Deep Layer (L5, L6) | (Lake BB *et al.* 2016) ^49^ | |
| *OXP2* | Deep Layer (L6) | (Lake BB *et al.* 2016) ^49^ | |

**Table S3. Antibodies and probes used in immunohistochemistry (IHC) and hybridization chain reaction – fluorescence in situ hybridization experiment (HCR-FISH).**

| **Primary and Secondary Antibodies for IHC** | | | | |
| --- | --- | --- | --- | --- |
| **Antibody** | **Host** | **Dilution** | **Manufacturer; Catalog number** | **RRID** |
| anti-MAP2 | Guinea pig | 1: 500 | Synaptic Systems; 188 004 | AB_2138181 |
| anti-m1 receptor | Goat | 1:500 | Abcam; ab77098 | AB_1523990 |
| anti-m3 receptor | Rabbit | 1:500 | Novus; NB100-58977 | AB_877677 |
| anti-guinea pig DyLight 405 | Donkey | 1:200 | Jackson Immunoresearch; 706-476-148 | AB_2632564 |
| anti-goat Alexa Fluor 750 | Donkey | 1:200 | Abcam; ab175745 | AB_2924800 |
| anti-rabbit Alexa Fluor 568 | Donkey | 1:200 | Invitrogen; A10042 | AB_2534017 |
| **Target Probe, Amplifier and Flurophore Combinations for HCR-FISH** | | | | |
| **Target** | **HCR Amplifier** | **Fluorophore** | | |
| *CHRM1* | B1 | Alexa Fluor 488 | | |
| *CHRM3* | B3 | Alexa Fluor 647 | | |

**Table S4. Between-*CHRM* subtype DEGs for pair-wise comparisons between *CHRM1+ vs CHRM2+ v CHRM3+* ExNs and InNs**

see excel file

**Table S5. Significantly enriched KEGG and GO Biological Processes Terms derived from DEGs between *CHRM1+ vs CHRM2+ v CHRM3+* ExNs and InNs**

see excel file

**Table S6. Between-Region DEGs and significantly enriched KEGG and GO Biological Processes Terms for ACC vs LPFC ExNs and InNs**

see excel file
